# A CPF-like phosphatase module links transcription termination to chromatin silencing

**DOI:** 10.1101/2023.07.06.547976

**Authors:** Eduardo Mateo-Bonmatí, Miguel Montez, Robert Maple, Marc Fiedler, Xiaofeng Fang, Gerhard Saalbach, Lori A Passmore, Caroline Dean

## Abstract

The interconnections between co-transcriptional regulation, chromatin environment and transcriptional output remain poorly understood. Here, we investigate the mechanism underlying RNA 3’ processing-mediated Polycomb silencing of Arabidopsis *FLOWERING LOCUS C* (*FLC*). We show a requirement for APRF1, a homologue of yeast Swd2 and human WDR82, known to regulate RNA Pol II during transcription termination. APRF1 interacts with TOPP4 (yeast Glc7/human PP1) and LD, the latter showing structural features found in Ref2/PNUTS; all components of the yeast and human phosphatase module of the CPF 3’end processing machinery. LD has been shown to co-associate *in vivo* with the histone H3 K4 demethylase FLD. We show APRF1 and LD couple CPF-mediated cleavage and polyadenylation with removal of H3K4 monomethylation in the body of *FLC*, and this influences subsequent transcription. This work shows how transcription termination can change the local chromatin environment to modulate transcription of Arabidopsis *FLC* and affect flowering time.

## INTRODUCTION

The relationship of chromatin and transcription in gene regulation is complex involving feedback mechanisms that make the interaction difficult to dissect. For example, chromatin influences RNA Polymerase II (RNA Pol II) processivity i.e., the likelihood of transcription reaching the end of the gene, which can lead to alternative splicing or early termination. In turn, these co-transcriptional steps feed-back to influence the chromatin state (Berry *et al*. 2015, Muniz *et al*. 2021, Zumer *et al*. 2021). One locus where this complexity has been studied in detail is Arabidopsis *FLC* (*FLOWERING LOCUS C*). *FLC* encodes a floral repressor and quantitative variation in *FLC* transcription has been an important determinant in adaptation of Arabidopsis accessions to a wide range of climates. For instance, high *FLC* expression upon germination in autumn enables an overwintering reproductive strategy with winter-induced Polycomb silencing of *FLC* aligning flowering with spring (Michaels and Amasino 1999). In contrast, low *FLC* expression through a developmentally induced Polycomb silencing leads to a rapid-cycling reproductive strategy, enabling multiple generations per year in some climates.

There has been extensive analysis of the components regulating *FLC* transcription to understand the developmentally induced silencing. The first repressor identified was FCA (FLOWERING CONTROL LOCUS A), an RRM-containing RNA binding protein (Macknight *et al*. 1997) that directly binds to *FLC* antisense transcript *COOLAIR* nascent transcripts, an association promoted by the presence of an R loop (Xu *et al*. 2021b). Forward genetic screens identified additional repressors that include other RNA binding proteins, RNA 3’ processing factors and chromatin modifiers (Wu *et al*. 2020). These factors have been found to promote proximal termination of both *COOLAIR* in seedlings and sense *FLC* transcripts in the embryo (Schon *et al*. 2021). Suppressor genetics and proteomic analysis showed that these co-transcriptional activities function through three factors that associate with each other *in vivo* - FLD (FLOWERING LOCUS D; a H3K4 demethylase)-LD (LUMINIDEPENDENS; TFIIS domain protein)-SDG26 (a SET domain protein) (Liu *et al*. 2007, Fang *et al*. 2020). These induce histone H3K4 demethylation across the locus to suppress *FLC*. SDG26 physically associates with FY/WDR33, a Cleavage and Polyadenylation Specificity Factor (CPSF; CPF in yeast) component after cross-linking (Fang *et al*. 2020), and both reduce H3K4me1 and H3K36me3 accumulation at *FLC*. This antagonizes transcription enabling a switch to PRC2-mediated H3K27me3 accumulation, so reducing transcriptional initiation and elongation rates (Wu et al 2016). The *FLC* silencing mechanism thus involves co-transcriptional processes that mediate chromatin modifications, which in turn feedback to reinforce the co-transcriptional processing. The factors involved are all evolutionarily conserved and affect many genes in Arabidopsis (Sonmez *et al*. 2011, Inagaki *et al*. 2021), raising the possibility that the mechanism may be broadly relevant in gene regulation.

A key question that remains is how proximal termination delivers a changed histone environment that enables the Polycomb Repressive Complex 2 (PRC2) switch. In the work described here we found a robust interaction of the FLD-LD-SDG26 complex with Arabidopsis APRF1 (ANTHESIS PROMOTING FACTOR 1), homologous to CPF phosphatase module component Swd2/WDR82. APRF1 also interacts with TOPP4 (TYPE ONE SERINE/THREONINE PROTEIN PHOSPHATASE 4), homologous to CPF phosphatase module component Glc7/PP1. The CPF phosphatase module dephosphorylates the RNA Pol II and its partners via the Glc7/PP1, an activity that promotes transcription termination through effects on RNA Pol II elongation/processivity (Schreieck *et al*. 2014, Carminati *et al*. 2023, Rodríguez-Molina *et al*. 2023). Close examination revealed that LD is structurally related to Ref2 in yeast and PNUTS in mammals. Ref2/PNUTS act in yeast/mammalian complexes as the regulatory subunit of the CPF phosphatase module. Using molecular and genetic tools we show that APRF1-dependent RNA processing activities function in the same co-transcriptional pathway as the FLD-LD-SDG26 chromatin modifier complex to promote proximal termination of the antisense *COOLAIR* transcripts, alter *FLC* chromatin environment, and affect *FLC* transcriptional output. This chromatin environment reinforces proximal termination choice so providing the molecular feedback necessary to stably maintain a low transcription state.

This work describes the mechanism linking transcription termination/ regulation of RNA Pol II and histone demethylation. APRF1, a structural component of a CPF-like phosphatase complex, directly links transcription termination with histone demethylase activity to alter the local chromatin environment and provide a mechanism resulting in graded repression of transcription. How this leads to the switch to Polycomb silencing was not resolved as none of the proteomic experiments identified Polycomb components. Our accompanying paper describes the use of mathematical modelling and experiments to elucidate how the mechanism described here sets the level of productive (processive) transcription that promotes the digital switch to the Polycomb silenced state (Menon *et al*. 2023).

## RESULTS

### The FLD complex robustly associates with APRF1, the homolog of yeast Swd2

We previously reported that the histone demethylase homolog FLD (FLOWERING LOCUS D) associates with LD (LUMINIDEPENDENS) and SDG26 (SET DOMAIN GROUP 26) *in vivo*; each tagged version of these three proteins enriched the other two partners in co-immunoprecipitation experiments (Fang *et al*. 2020). Interestingly, each of these proteins also co-immunoprecipitated with ANTHESIS PROMOTING FACTOR1 (APRF1; Fang *et al*. 2020), a result recently confirmed in an independent analysis (Qi *et al*. 2022).

APRF1 is a WD40-repeat protein encoded by At5g14530. We obtained a T-DNA insertion line [WiscDsLox_489; *aprf1-9*; Kapolas *et al*. (2016)] and analysed the flowering time and *FLC* expression. Both *FLC* spliced and unspliced transcripts were significantly upregulated in the mutant line, and accordingly, *aprf1-9* plants were late flowering (Figure 1A-C). The insertion interrupts the 9^th^ of 10 exons, so it was possible that this mutant retained some APRF1 function. To address this possibility, we designed a CRISPR-Cas9 transgene to generate a full knock-out for *APRF1*. Using a sgRNA targeting the second exon and screening for edited plants, we found transgene-free T_2_ plants carrying a 5-nt deletion which creates an in-frame premature stop codon (Figure S1). This new allele was named *aprf1-10* and was as late flowering as *aprf1-9* (Figure 1B). We crossed *aprf1-9* to *aprf1-10* and analysed *FLC* expression levels of the single and heterozygous mutants. All had similarly upregulated *FLC,* confirming their allelism and the role of APRF1 on *FLC* repression (Figure 1C).

**Figure 1.**
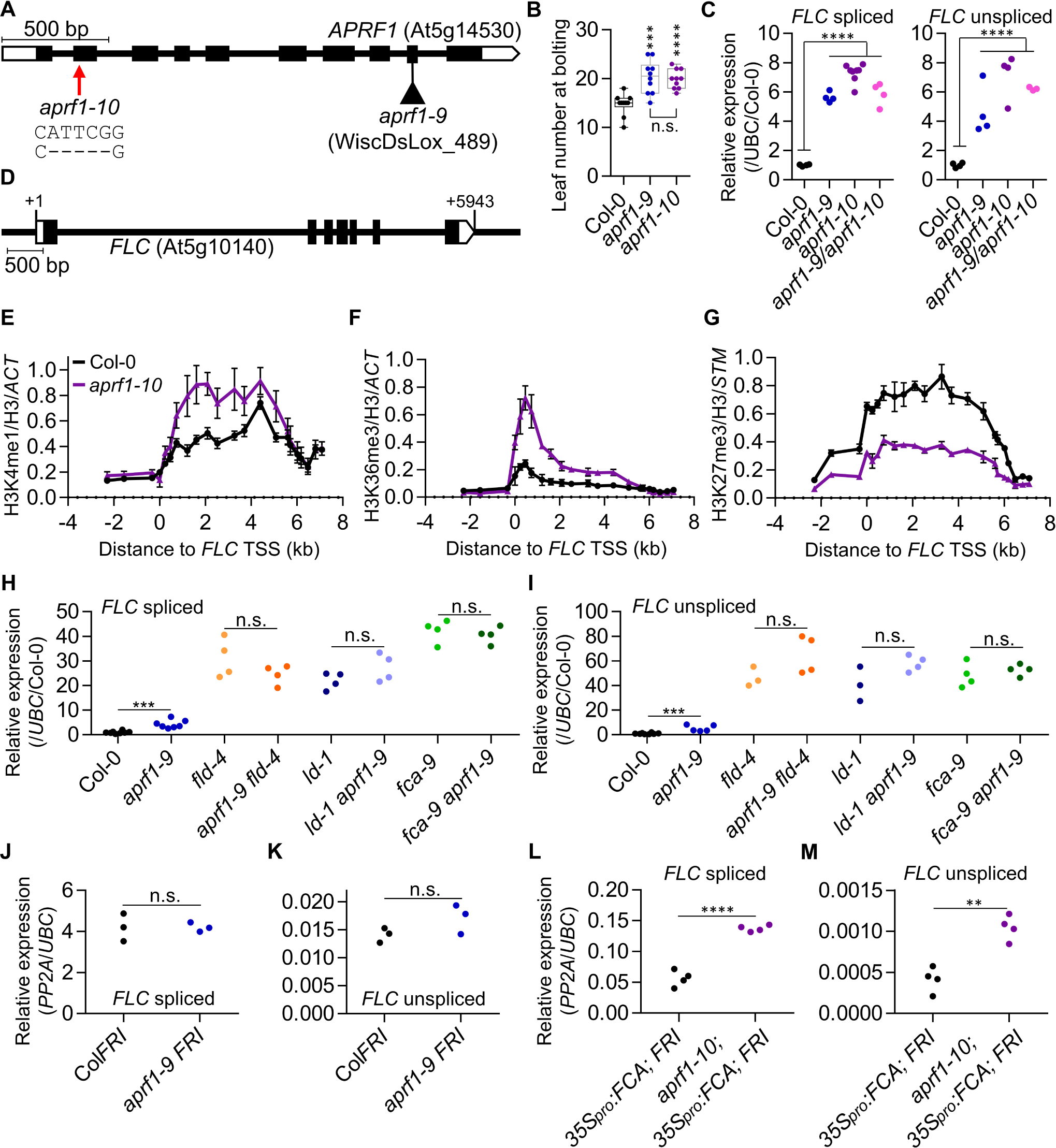
APRF1, a robust interactor of the FLD complex, functions genetically in the FCA pathway. (**A**) Architecture of the *APRF1* gene and illustration of the nature and location of the mutations studied in this work. Boxes indicate exons and lines introns. White boxes represent untranslated regions. The triangle represents a T-DNA insertion (*aprf1-9*); the red arrow points to the location of the CRISPR/Cas9-derived deletion (*aprf1-10*). (**B**) Boxplot representing the leaf number at bolting of the wild-type Col-0 and both *aprf1* mutants. Each dot represents the score of a single plant. (**C**) Relative values of *FLC* spliced (left) and unspliced (right) in the wild-type Col-0, both *aprf1* mutants and F_1_ *aprf1-9/aprf1-10* hybrid plants. Values were normalized to the housekeeping *UBC* gene and to Col-0. (**D**) Schematic diagram showing *FLC* gene structure following the guidelines described for (**A**). +1 indicates the transcriptional start site (TSS). (**E-G**) ChIP analysis of H3K4me1 (**E**), H3K36me3 (**F**), and H3K27me3 (**G**) levels at *FLC* in Col-0 and *aprf1-10*. Numbers in x axis represent the distance in kilobases to the *FLC* TSS and numbers in the y axis correspond to relative enrichment of the corresponding histone mark. Each dot represents an amplicon. Values were normalized to H3 and to *ACT7* (for H3K4me1 and H3K36me3) or *STM* (for H3K27me3). (**H-M**) Relative values of *FLC* spliced (**H, J, L**), and unspliced (**I, K, M**) in various genetic backgrounds. Values were normalized to the housekeeping gene *UBC* (**H-M**) and Col-0 (**H, I**) or *PP2A* (**J-M**). Asterisks indicate statistically significant differences to Col-0 (**B, C, H, I**), and to C2 (*35S_pro_:FCA*_γ_*; FRI*) (**L, M**), in a two-way Student’s *t* test (** *p* < 0.01, *** *p* < 0.001, and **** *p* < 0.0001). N.s. stands for not statistically different. Scale bars indicate 500 bp (**A, D**). Experiments were performed using 2-week-old seedlings grown in long days conditions with n ≥ 3 (**C, E-M**).

### Swd2 function is functionally diverged in Arabidopsis

APRF1 is one of the two Arabidopsis orthologs of the *Saccharomyces cerevisiae* Swd2 [(Figure S2A)(Fiorucci *et al*. 2019)]. Swd2 has been found to play a role in two very different complexes. One of which being the COMplex of Proteins Associated with Set1 (COMPASS), responsible for the co-transcriptional deposition of H3K4me3 where Swd2 promotes the interaction between Set1 and the RNA Pol II Carboxy Terminal Domain (CTD) (Bae *et al*. 2020). Additionally, Swd2 has been found as a component of the phosphatase module of the Cleavage and Polyadenylation Factor (CPF; CPSF in higher eukaryotes) and the Associated with Pta1 (APT) complexes, which signal transcriptional termination (Casanal *et al*. 2017, Lidschreiber *et al*. 2018). These apparently opposite roles on transcription in yeast motivated us to test if the Arabidopsis orthologs *APRF1* (also known as *Swd2-like A/S2LA*) and *S2LB* had functionally diverged. *FLC* spliced and unspliced levels were assayed in an insertional allele of *S2LB* (Figure S2B) and the double mutant *aprf1-9 s2lb.* Opposite to *aprf1-9*, *s2lb* showed a significant reduction in *FLC* expression, while the double mutant had *FLC* levels indistinguishable from the *aprf1-9* single mutant (Figure S2C). We further analysed the effects on *FLC* expression in a representative mutant of the COMPASS activity, *atx1-2,* an insertion allele in *ARABIDOPSIS TRITHORAX1,* which encodes one of the methyltransferases. In line with the results observed for *s2lb*, both *FLC* spliced and unspliced were significantly downregulated in *atx1-2* compared to Col-0 (Figure S2D). These results perfectly match with previous work reporting *FLC* downregulation in other COMPASS mutants defective in components such as WDR5 (Jiang *et al*. 2009), RBL, or ASH2R (Jiang *et al*. 2011). FLD-mediated repression of *FLC* requires H3K4me1 removal (Fang *et al*. 2020, Inagaki *et al*. 2021). Considering the tight link between the FLD complex and APRF1, we performed chromatin immunoprecipitation coupled with quantitative PCR (ChIP-qPCR) to quantify H3K4me1 at *FLC*. We found higher levels of H3K4me1 across the locus in *aprf1-10* compared to Col-0 (Figure 1E), matching results in *ld-1* and *fld-4* (Figure S2E; Fang *et al*. 2020) or genome-wide (Inagaki *et al*. 2021). In contrast, *s2lb* plants showed slightly lower levels of H3K4me1 than the Col-0 (already low), in line with COMPASS dysfunction (Figure S2E). The contrasting phenotypes shown by *s2lb* and *aprf1* suggest that in Arabidopsis, the ancestral Swd2 may have sub-functionalized, with S2LB working through the COMPASS complex and therefore activating *FLC,* and APRF1 working with the 3’ processing machinery to repress the locus (Figure S2F).

H3K4me1 binds SDG8 (Liu and Huang 2018), a histone methyltransferase that deposits H3K36me3, an active histone modification, which is mutually exclusive to the repressive PRC2-deposited H3K27me3 at *FLC* (Yang *et al*. 2014). Consistent with the *FLC* upregulation and the increased H3K4me1 levels in *aprf1-10*, ChIP-qPCR analyses also showed that H3K36me3 and H3K27me3 were upregulated and downregulated, respectively. Thus, APRF1 has a role in establishing a silent chromatin state at *FLC* (Figure 1F, G).

### APRF1 functions downstream FCA

Working genetically upstream of FLD functionality, FCA promotes proximal termination on both strands of *FLC* (Simpson *et al*. 2003, Schon *et al*. 2021). To obtain genetic evidence of the relationship between *APRF1* and *FLD, LD* or *FCA*, we crossed *aprf1-9* with mutants in subunits of the FLD complex such as *fld-4* or *ld-1*, and the core component of the pathway *fca-9*. *FLC* levels in the double mutants revealed an epistatic relationship between *FLD, LD* or *FCA* and *APRF1* (Figure 1H, I), further demonstrating the genetic connection between *APRF1* and the *FLC* repression machinery. To study the effects of loss of *APRF1* in a high transcriptional environment, but without perturbing the FCA pathway, we introgressed a functional *FRIGIDA* (*FRI*) (Johanson *et al*. 2000) into *aprf1-9*. FRI functions as an *FLC* transcriptional activator, and like other anti-terminators (Gregersen *et al*. 2019), promotes distal polyadenylation of both *FLC* and *COOLAIR*, thus antagonizing the co-transcriptional repression mechanism (Schon *et al*. 2021). Levels of *FLC* in *aprf1-9 FRI* were the same as Col*FRI,* likely due to an overwriting effect of *FRI* compared to the relatively modest *FLC* upregulation of *aprf1-9* (Figure 1J, K). Finally, we previously generated a sensitised transgenic system called C2, in which the chromatin of *FLC* is silenced even in the presence of an active *FRI* by the overexpression of the *FCA* through the transgene *35S_pro_:FCA*_γ_ (Liu *et al*. 2010, Wu *et al*. 2020). Mutations affecting FCA-downstream processes compromise the FCA-mediated *FLC* chromatin silencing. Introgression of *aprf1-10* into the C2 background resulted in significant release of *FLC* repression, demonstrating a role for APRF1 downstream of FCA (Figure 1L, M).

### APRF1 reciprocally interacts with LD, the plant homolog of the phosphatase regulatory subunit Ref2/PNUTS

To further characterize the role of APRF1 on *FLC* repression we performed CrossLinked Nuclear ImmunoPrecipitation and Mass Spectrometry (CLNIP-MS) using 10-days-old seedlings of a FLAG-tagged version of APRF1 (Qi *et al*. 2022). We found that the top-hit was LD, thus confirming their reciprocal interaction (Figure 2A; Table S1). Consistent with ours (Fang *et al*. 2020) and other reports (Qi *et al*. 2022) among the top-hits we also found FLD (Figure 2A). Interestingly, one of the highest hits was the histone H2A.W.7, a variant exclusively found on constitutive heterochromatin (Lorković *et al*. 2017, Jamge *et al*. 2023), suggesting a generic role for APRF1 in co-transcriptional gene repression. Given this robust APRF1-LD interaction, we considered if LD could be the homolog of one of the yeast Swd2 partners in the CPF or the APT phosphatase modules (Casanal *et al*. 2017, Lidschreiber *et al*. 2018). Ref2 was an interesting candidate since both Ref2 and LD are highly unstructured proteins (Figure 2B). Ref2 is key for the interaction between CPF and the RNA Pol II and is a phosphatase regulatory subunit, providing substrate specificity to the phosphatase Glc7 (Russnak *et al*. 1995, Nedea *et al*. 2008, Carminati *et al*. 2023). Ref2, and its putative metazoan ortholog PNUTS, are largely disordered proteins apart from a TFIIS motif at the N terminal region of the protein which consists of a compact four-helix bundle (Figure 2B). Intriguingly, despite its apparent key role in an otherwise highly conserved 3’ processing machinery, Ref2 shows no obvious homology with an Arabidopsis protein. We then performed BLAST searches to find potential orthologs in Arabidopsis for PNUTS (Allen *et al*. 1998). BLAST algorithms showed the best hit for PNUTS in Arabidopsis is LD (Figure S3). We aligned the sequences of Ref2, PNUTS and LD, and although the degree of conservation is low (Figure S4), improved slightly when aligning only LD and PNUTS (Figure S5). Nevertheless, all of them share similar features, a TFIIS domain in the N-terminus in an overall highly unstructured protein (Figure 2B, S4B-D). PNUTS is known to interact with WDR82, a bona-fide homolog of APRF1 (Figure S6), and provides substrate specificity to PP1 phosphatases, like Glc7. We then searched for phosphatases among the APRF1 interactors, finding a highly significant interaction with TYPE ONE SERINE/THREONINE PROTEIN PHOSPHATASE4 (TOPP4; Figure 2A), a protein with very high homology to Glc7 and PP1 (Figure S7) and proven phosphatase activity *in vivo* (Wang et al. 2022), and C-TERMINAL DOMAIN PHOSPHATASE-LIKE 3 (CPL3), homolog to yeast FCP1 whose role activating FRIGIDA complex activity has been reported (Shen *et al*. 2022). We have experimentally validated the interaction between APRF1 and TOPP4 *in planta* using transient co-immunoprecipitation (co-IP) in *Nicotiana benthamiana* leaves of *TOPP4_pro_:TOPP4-3−FLAG* and *APRF1_pro_:APRF1-mVENUS* (Figure 2C). The IP-MS experiment also found a significant interaction between APRF1 and CPSF100, a structural component of the CPSF homolog of the yeast Cft2. In summary, our results suggest that APRF1/LD/TOPP4 form a plant equivalent to the yeast (Swd2/Ref2/Glc7) and human (WDR82/PNUTS/PP1) CPF phosphatase modules.

**Figure 2.**
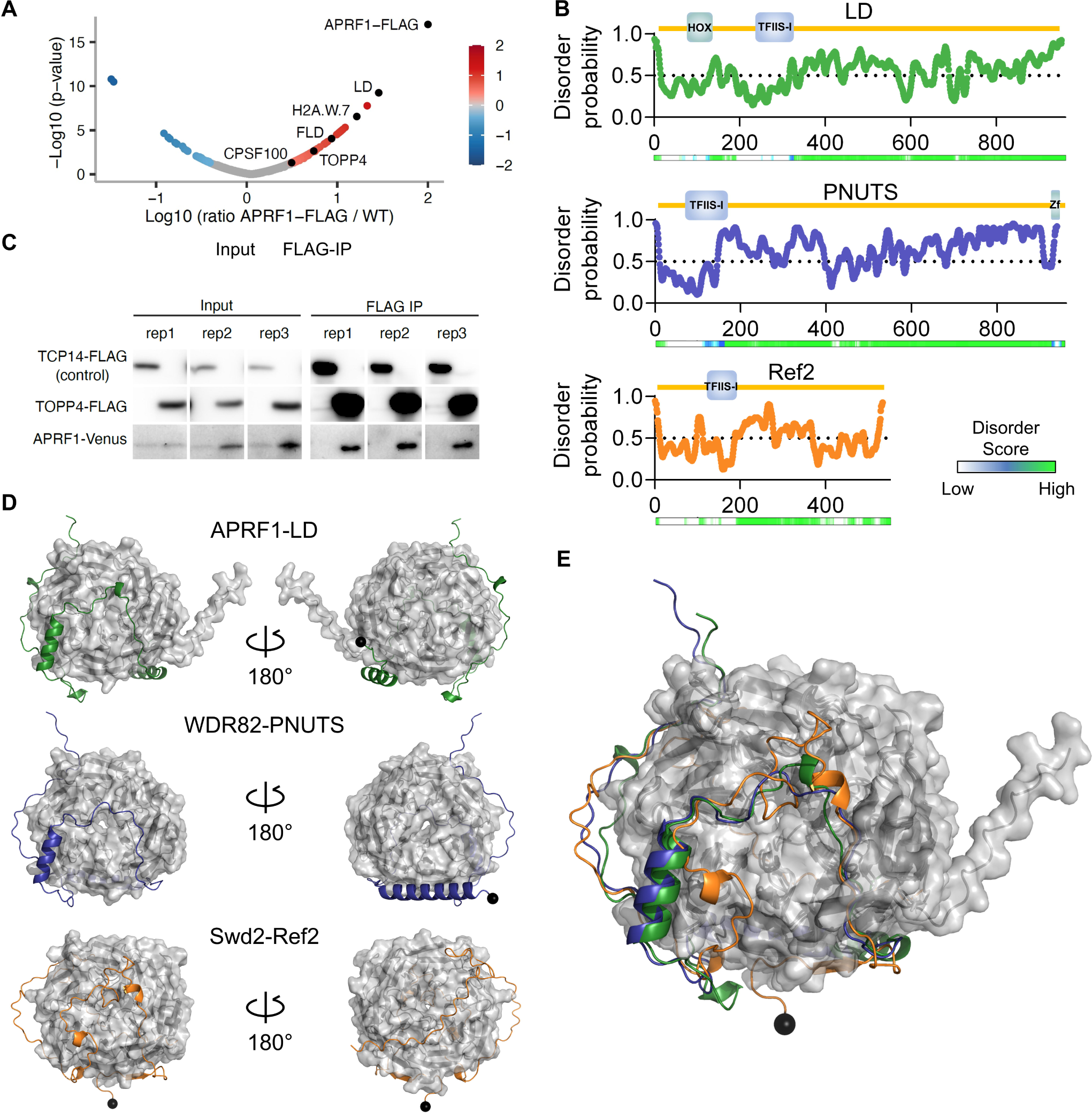
LD-APRF1-TOPP4 form a plant CPSF-like phosphatase module. (**A**) Volcano plot showing the relative protein abundance in log-10 scale ratio of immunoprecipitated samples from APRF1-3xFLAG to control Col-0 samples. Red dots highlight proteins enriched in the APRF1-3xFLAG samples. APRF1-FLAG, LD, H2A.W.7, FLD, TOPP4, and CPSF100 are shown as black dots. (**B**) Schematic representation of protein size, annotated domains, disorder probability, and disorder score for LD, PNUTS (*Homo sapiens*), and Ref2 (*Saccharomyces cerevisiae*). Individual amino acid score for disorder probability were obtained with the online Protein DisOrder prediction System (PrDOS) and plotted using Graphpad. Disorder scores were obtained by D^2^P^2^ (Oates *et al*. 2013) and shown in a colour scale. (**C**) Co-immunoprecipitation results obtained in transiently transform leaves of *N. benthamiana* with APRF1-mVENUS, TOPP4-3xFLAG, and the control line TCP14-FLAG. Three replicates of the same experiment are shown (**D**) AlphaFold2 predictions of complexes between APRF1-LD, WDR82-PNUTS, and Swd2-Ref2. WD40 orthologs are shown in grey surface representation, whereas TFIIS orthologs are shown in cartoon; the N-terminal amino acid of the predicted TFIIS proteins are denoted by black spheres. (**E**) Overlay of the three predictions shown in (**D**).

Further exploring these parallels, we carried out an *in-silico* prediction using AlphaFold2 of the interaction between APRF1-LD and their yeast and human counterparts (Figure 2D). Despite the low sequence homology, predictions support conservation of structural features and contact points between APRF1/Swd2/WDR82 and LD/Ref2/PNUTS thus supporting their functional equivalence (Figure 2E). Taken together, the robust immunoprecipitation of LD and TOPP4 by APRF1 and the structural parallels between LD, Ref2, and PNUTS suggest that LD, APRF1, and TOPP4 form a CPF or CPF-like phosphatase module in Arabidopsis.

### FLD complex and RNA Pol II co-occupy *FLC* chromatin independently of FCA function

Available genome-wide data shows FLD binding correlates with actively transcribed genes and elongating RNA Pol II CTD phosphorylated at Ser2 or Ser5 (Inagaki *et al*. 2021). Similarly, the Arabidopsis FLD paralog LDL3 has been recently reported to work co-transcriptionally to remove H3K4me2 (Mori *et al*. 2023). RNA Pol II occupancy in the Arabidopsis genome often shows peaks near transcription termination sites (TTS), potentially linked to slow co-transcriptional termination events (Wu *et al*. 2016, Zhou *et al*. 2023). We found this was the case for *FLC* in Col*FRI* vs Col-0 (Figure 3A) and has been shown to be the case in *fca-9* or *fld-4* (Wu *et al*. 2016). To address co-occupancy with FLD, we performed ChIP-qPCR using a transgenic FLAG-tagged FLD with and without the transcriptional activator FRI, and non-transgenic control plants. Agreeing with the reported data (Inagaki *et al*. 2021), FLAG-FLD showed high enrichment at the 3’ region of *FLC* in a *FRI* genotype compared to Col-0 (*fri)*, with the latter close to the background signal (Figure 3B). To rule out the possibility that this enrichment was an effect of the transcriptional activator FRI and not the high transcription itself, we introgressed the FLAG-FLD transgene into the *fca-9* background. FLD enrichment at the 3’end of *FLC* in *fca-9 FLAG-FLD* was indistinguishable from *FRI FLAG-FLD* confirming FLD enrichment primarily associated to transcriptional activity (Figure 3B). *FLD* functions genetically downstream of *FCA* (Liu *et al*. 2007) so it was interesting that FLD-Pol II co-occupancy association was not affected by *fca-9*. Thus, even when FLD is located at *FLC*, it cannot function properly without FCA-mediated 3’ processing of the nascent transcript.

**Figure 3.**
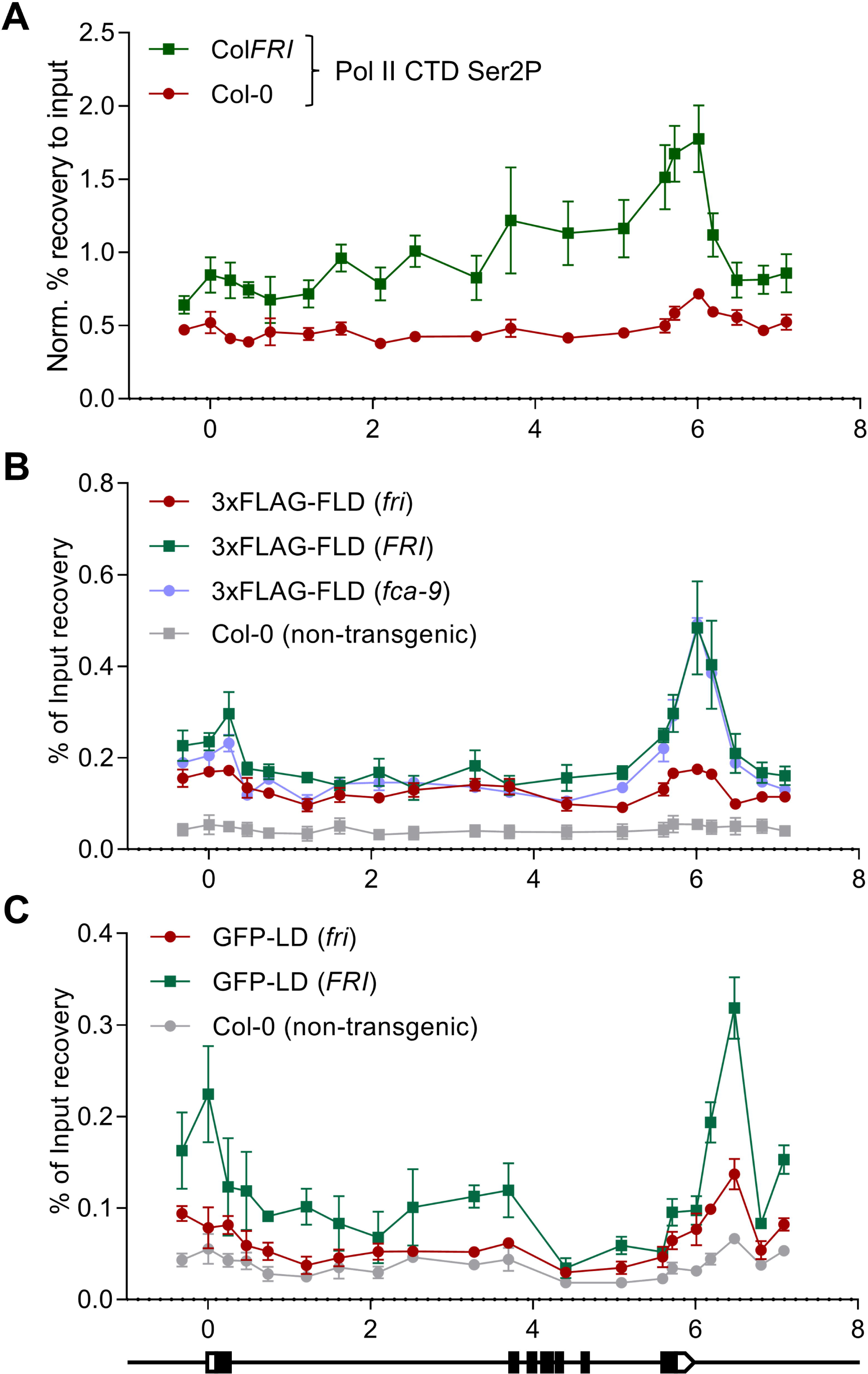
ChIP-qPCR co-occupancy profiles indicate that FLD and LD work co-transcriptionally to control *FLC* transcription. (**A**) Elongating RNA Pol II (Ser2P) ChIP profiles over *FLC* in a high (Col*FRI*) and low (*fri* or Col-0) transcriptional background. (**B**) ChIP binding profile over *FLC* of the *fld-4 FLD_pro_:3xFLAG-FLD* transgenic plants in different genetic backgrounds. (**C**) ChIP binding profile over *FLC* of the *ld-1 LD_pro_:GFP-LD* transgenic plants in different genetic backgrounds. *FLC* gene structure following the indications for Figure 1D. All the experiments were done with 2-week-old seedlings. Dots and error bars represent average ± s.e.m. of three replicates. (**A**) Results expressed in % recovery to input values normalized to the promoter of the housekeeping gene *ACT7* as in Mikulski *et al*. 2022. Results in (**B, C**) are expressed in % recovery to input.

We introgressed a GFP-tagged LD (Fang *et al*. 2020) into Col*FRI* to create a line with high *FLC* transcription and performed ChIP-qPCR experiments. LD enrichment, peaking at the *FLC* TTS, was only detected in the line where *FLC* transcription is high (Figure 3C). The shared enrichment of elongating Pol II, FLD- and LD is consistent with an *in vivo* association of the transcriptional machinery and the FLD complex as suggested by genome-wide data (Inagaki *et al*. 2021). Thus, we propose that FLD and LD associate with RNA Pol II, and therefore work co-transcriptionally, fitting with LD working as a CPF component equivalent to Ref2 or PNUTS.

### Inefficient termination in *aprf1* mutants leads to transcriptional read-through

Previous work on the FCA pathway found that most of the factors involved directly affected *FLC* antisense transcripts (*COOLAIR*) (Wu *et al*. 2020). To pursue a potential role of APRF1 in transcriptional termination, we analysed the transcription and processing of *COOLAIR*. In contrast to *FLC* levels which did not show any significant difference (Figure 1J, K), total *COOLAIR* levels were upregulated in double mutants containing *aprf1-9* (Figure 4A, B). This was particularly striking for the *aprf1-9 FRI* combination. *COOLAIR* transcripts are polyadenylated at many sites with major clusters at proximal sites (Class I) and distal sites coincident with the *FLC* promoter (Class II; Figure 4A) (Liu *et al*. 2010). An increase in proximal *COOLAIR* (Figure 4C) and no change in distal *COOLAIR* were found in *aprf1-9 FRI* (Figure 4D), with the proximal / distal ratio increased in *aprf1-9* double mutant combinations compared to single mutants (Figure 4E), unlike other mutations affecting the FCA pathway (Xu *et al*. 2021a). However, as with the *Arabis alpina FLC* ortholog (*PEP1*) (Castaings *et al*. 2014), we detect low abundance spliced *COOLAIR* transcripts polyadenylated around a medial site (Liu and Dean, unpublished), which we term *COOLAIR* Class III (Figure 4A). This isoform was significantly up-regulated specifically in *aprf1-9 FRI* (Figure 4F), with one specific spliced variant *COOLAIR* Class III.3 becoming the most abundant isoform, enriched in high transcription situations such as *fca-9* or Col*FRI* (Figure 4G, S9A, B). In line with the proposed divergent roles and consistent with the *FLC* sense expression profile (Figure S2C, D), *s2lb* and *atx1-2* mutants showed significantly lower levels of all the *COOLAIR* isoforms compared to Col-0 (Figure S9C). The high levels of the *COOLAIR* class III isoform seemed likely to reflect inefficient polyadenylation/transcript termination at the proximal site. To analyse the polyadenylation site choice in an unbiased and strand-specific manner in these two genotypes, we carried out a Quant-seq analysis of Col*FRI* and *aprf1-9 FRI*, to detect mRNA 3’-end. There were only a small number of reads on the *FLC* (sense) strand consistent with transcriptional readthrough (Figure S10). However, for *COOLAIR* the differences were large, with many reads indicating use of the medial poly A site (class III), distal *COOLAIR* readthrough, and alternative *COOLAIR* transcriptional starts (Figure 4H, S10). To determine if the effects were specific to loss of APRF1, we performed Quant-seq analysis of *fld-4* compared to the wild-type Col-0 (Figure S11). *FLC* sense transcripts were qualitatively the same, just quantitatively up-regulated in *fld-4* (Figure S11A, B). The few reads corresponding to *COOLAIR* were insufficient to make any conclusions (Figure S11B) so a library enrichment was performed using baits encompassing the 20-kb *FLC* genomic region (see Methods). As a control, we also performed this bait enrichment on the Col*FRI, aprf1-9 FRI* libraries (Figure S12). No medial poly A site (class III) were found in Col-0 or *fld-4* (Figure S11C, D), suggesting the termination defects are APRF1-specific and that the FLD downstream function is not involved in the termination process.

**Figure 4.**
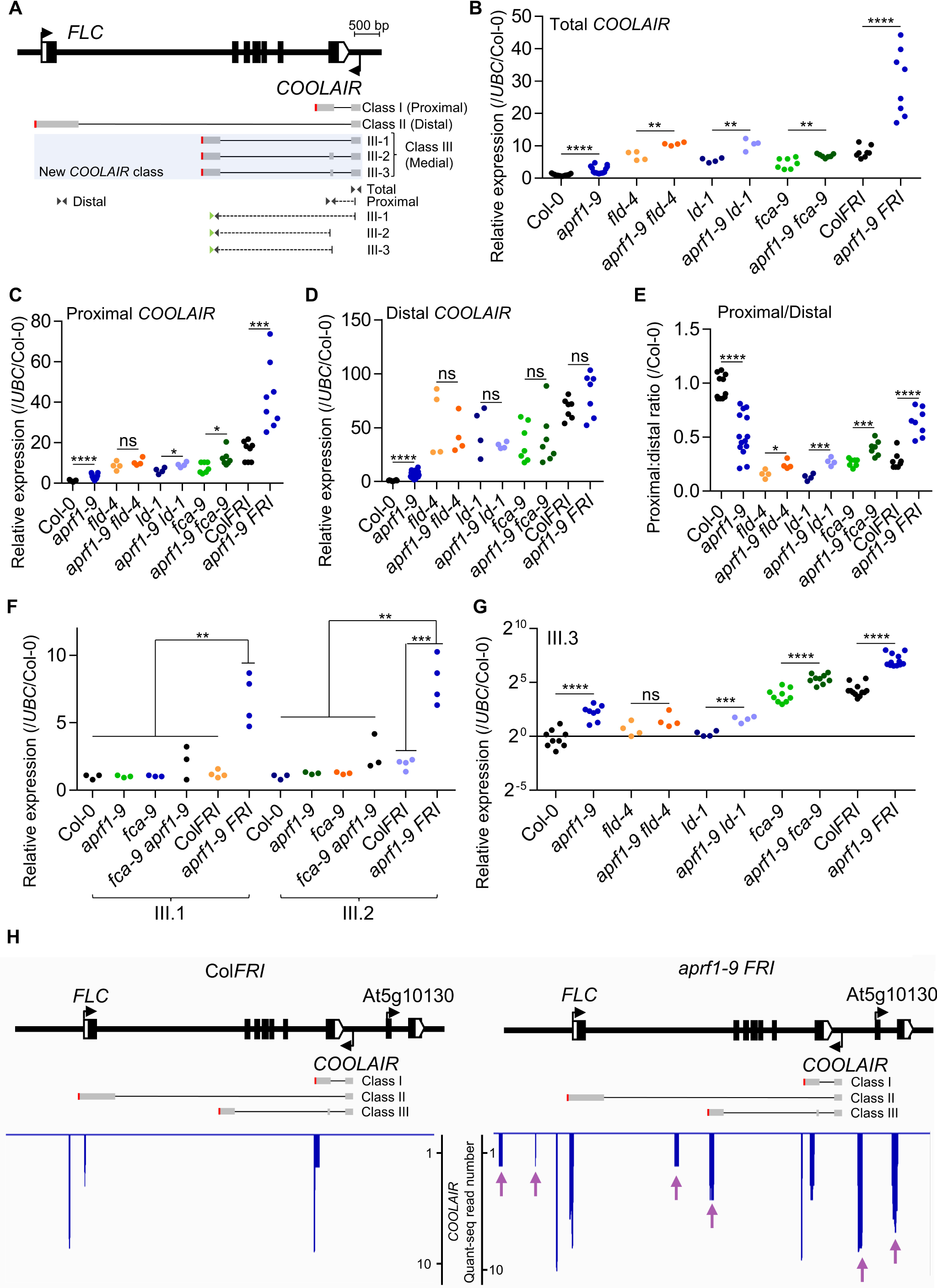
Mutations in *APRF1* trigger *COOLAIR* upregulation and an increase in a new medially polyadenylated *COOLAIR* isoform. (**A**) *FLC* architecture following the representation of Figure 1D with indication of different *COOLAIR* isoforms in grey. For simplicity, Classes I and II (proximal and distal) are each represented by one isoform. The new *COOLAIR* class III isoforms are highlighted in pale blue. Triangles represent primer pairs (not drawn to scale) used to measure the relative abundance by qRT-PCR. Dashed lines indicate primers spanning two *COOLAIR* exons. Green triangles indicate the primer used for *COOLAIR* class III retro-transcription. Red vertical lines represent polyadenylation sites. Scale bar indicates 500 bp. (**B-D**) Relative expression analyses of (**B**) total *COOLAIR,* (**C**) proximal *COOLAIR*, and (**D**) distal *COOLAIR* in different genotypes. (**E**) *COOLAIR* proximal-to-distal ratio in different genotypes. (**F, G**) Relative expression analyses of *COOLAIR* class (**F**) III.1 and III.2, and (**G**) III.3. Each dot represents a biological replicate analysed in triplicate. Expression values were normalized to the *UBC* gene and to Col-0. Asterisks indicate statistically significant differences to the indicated genotypes in a Student’s *t* test (* *p* < 0.05, ** *p* < 0.01, *** *p* < 0.001, and **** *p* < 0.0001). N.s. stands for not statistically different. All the experiments were performed in 2-week-old seedlings grown in long days conditions. (**H**) *COOLAIR* strand Quant-seq results of Col*FRI* and *aprf1-9 FRI* seedlings. *FLC* locus is represented as in panel (**A**). The blue spikes indicate reads supporting a polyadenylation site. Purple arrows point to clusters of reads present in *aprf1-9 FRI* but absent in Col*FRI*.

Both Quant-seq datasets revealed more than two hundred commonly misregulated genes in *aprf1-9 FRI* and *fld-4* compared to the corresponding wild-type strain (Figure S13A, B; Table S2, S3). Among the genes upregulated in *fld-4* and even more on those commonly upregulated in both mutant backgrounds, we observed a generalized shift from proximal to distal polyadenylation site choice in *aprf1-9 FRI* compared to Col*FRI* (Figure S13C-E). We also found Quant-seq signals compatible with read-through or inefficient proximal polyadenylation in other genes (Figure S13F, G). Taken together, our Quant-seq analyses suggest *APRF1* loss leads to inefficient transcriptional termination at many loci in the Arabidopsis genome.

To further investigate co-transcriptional changes we analysed chromatin-bound RNA (chRNA) of Col-0, *aprf1-9*, Col*FRI* and *aprf1-9 FRI* at *FLC*. *FLC* (sense) chRNA levels of *aprf1-9* were higher than Col-0, as *aprf1-9* compromises *FLC* repression. PCR amplicons for sense *FLC* levels in Col*FRI* and *aprf1-9 FRI* gave similar values at the 5’ and 3’ ends of the locus, but there were differences in the central region, pointing to complex effects of APRF1 on *FLC* Pol II processivity (Figure 5A). However, the most striking differences were on the *COOLAIR* strand (antisense) (Figure 5A). RNA was detected in regions corresponding to *COOLAIR* class I and class II in Col*FRI*, but in *aprf1-9* genotypes (*fri* and *FRI*) this signal extended to regions that cover both *COOLAIR* Class I and Class III transcripts. This likely represents antisense transcriptional readthrough, supporting a role for APRF1 as part of the RNA Pol II termination machinery (Figure 5A).

**Figure 5.**
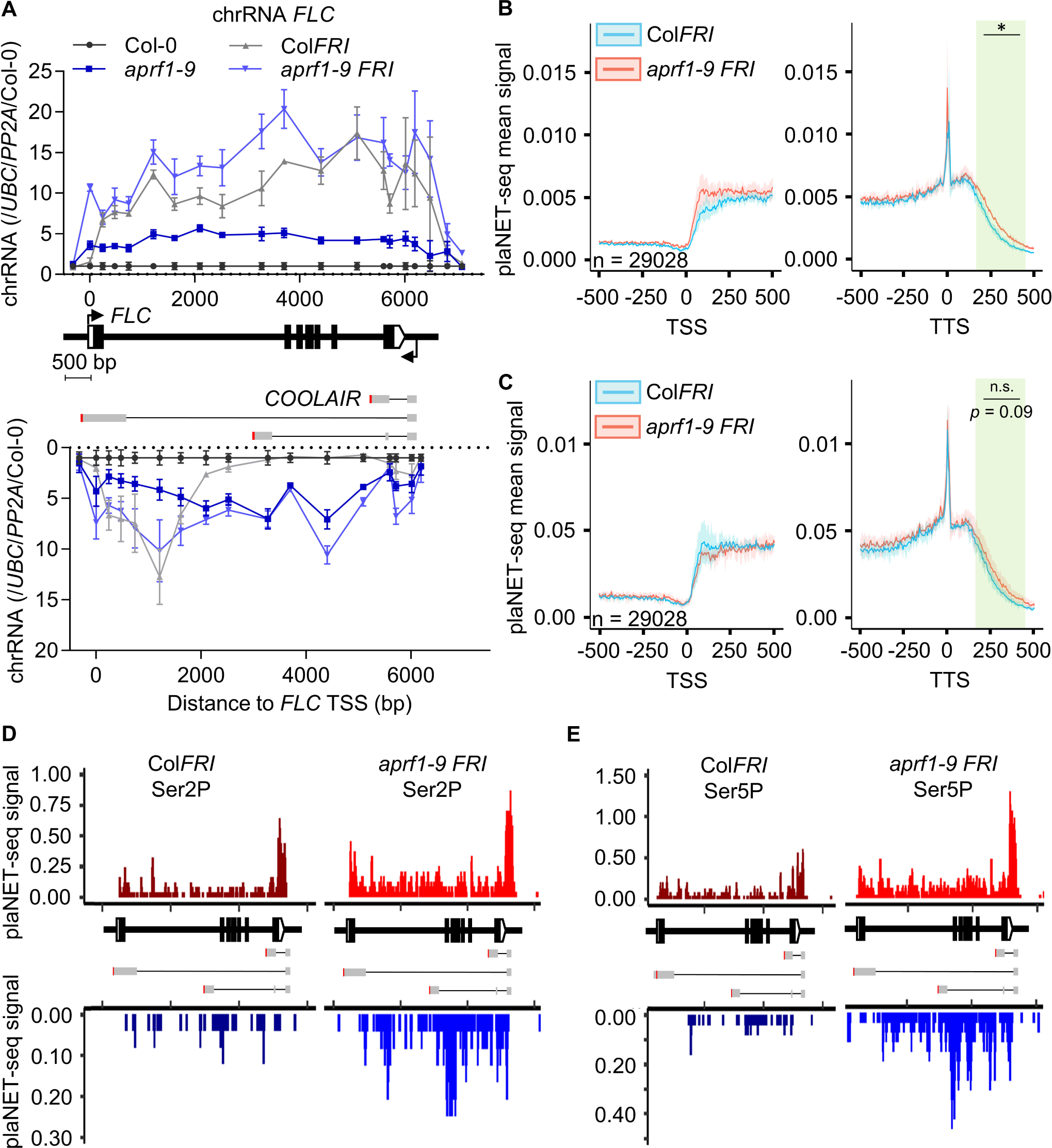
*COOLAIR* transcriptional readthrough correlates with changed phosphorylation of the RNA Pol II carboxy terminal domain. (**A**) Chromatin-bound RNA levels of *FLC* and *COOLAIR* in *aprf1-9* mutants with and without functional *FRI*. Data was normalized to *UBC* and *PP2A* and is shown as fold-change to wildtype Col-0. Dots correspond to amplicons of the *FLC* (upper chart) and *COOLAIR* (bottom chart) transcripts, represented as average ± s.e.m. of three biological replicates. (**B**, **C**) plaNET-seq metaplots at TSS and TTS of Col*FRI* and *aprf1-9 FRI* using Ser2P (**B**) and Ser5P (**C**) antibodies. Asterisk indicates statistically significant differences between *aprf1-9 FRI* and Col*FRI* in a one-way ANOVA with multiple comparisons. (**D**, **E**) plaNET-seq profiles over the *FLC* locus of Col*FRI* and *aprf11-9 FRI* using Ser2P (**D**) and Ser5P (**E**) antibodies. Upper and bottom charts show plaNET-seq profiles corresponding to *FLC* and *COOLAIR* strands, respectively.

### APRF1 affects RNA Pol II CTD Ser2/5 phosphorylation

Different mechanistic models, not mutually exclusive, have been proposed for transcription termination (Mo *et al*. 2021, Rodríguez-Molina *et al*. 2023). Current thinking based on studies in different organisms is that CPF-RNA Pol II recognition of polyadenylation site (PAS) triggers the nascent RNA cleavage. This generates a free 5’ end on the cleaved RNA which is a substrate for 5’-3’ ribonucleases such as XRNs (5’-3’ EXORIBONUCLEASEs) that degrade the nascent transcript and dislodge RNA Pol II from the chromatin (Cortazar *et al*. 2019, Rodríguez-Molina *et al*. 2023). Transcription of the PAS also triggers a conformational change in RNA Pol II, potentially driven by dephosphorylation of the CTD and/or co-factors, which slows-down RNA Pol II. To ascertain if *aprf1* mutants affect RNA Pol II CTD modifications, we carried out ChIP-qPCR experiments using antibodies targeting the RNA Pol II CTD phosphorylated residues Ser2, Ser5 and Tyr1 comparing *aprf1-10* with Col-0. The phosphorylated RNA Pol II was detected at higher levels in *aprf1-10,* consistent with the locus being more actively transcribed (Figure S14A). We repeated our analyses comparing *ld-1* and the double mutant *aprf1-9 ld-1* given that both genotypes have similar *FLC* RNA levels (Figure 1H). RNA Pol II Ser2P and Tyr1P levels were higher than Col-0 but identical between the single and the double mutant (Figure S14B). However, Ser5P levels were higher in the *ld-1 aprf1-9* plants, indicating a role for APRF1 on RNA Pol II CTD Ser5 dephosphorylation.

Since our previous analyses revealed levels of *COOLAIR* class III (Figure 4F-H) and transcriptional readthrough (Figure 5A) were particularly clear in a *FRI* background without affecting the overall *FLC* expression levels (Figure 1J, K), we also performed a ChIP-qPCR experiments using the RNA Pol II CTD antibodies for Col*FRI* and *aprf1-9 FRI*. RNA Pol II Ser2P and Tyr1P levels were identical, while there was a slight increase in RNA Pol II Ser5P towards the 5’ end of the locus (Figure S14C). In order to obtain genome-wide, more sensitive, and strand-specific information on the effects of *APRF1* in this background, we carried out plant Native Elongating Transcript sequencing experiments (plaNET-seq) (Kindgren *et al*. 2020) using the native CTD phosphorylated residues Ser2P and Ser5P. We observed a statistically significant generalized readthrough in *aprf1-9 FRI* using Ser2P and differences using Ser5P which were not significant (Figure 5B, C). At the *FLC* locus, and in striking contrast to the low-resolution and not strand-specific ChIP experiments, plaNET-seq revealed an increase in Ser2P in *aprf1-9 FRI* in the (*FLC*) sense strand compared to Col*FRI*, with a higher peak at the 3’ region of the locus, in agreement with *aprf1-10* ChIP results (Figure 5D). The *COOLAIR* strand showed greater differences, with *aprf1-9 FRI* having higher levels overall than Col*FRI* and a sharp accumulation close to the class III polyadenylation site (Figure 5D). Ser5 phosphorylation differences were even bigger, with an obvious accumulation on both strands and increased Ser5P near the class III polyadenylation site (Figure 5E). These results could suggest a role for the APRF1-termination complex in removal of RNA Pol II Ser2P/Ser5P at *FLC.* However, we cannot rule out the possibility that the CTD hyperphosphorylation we observe is an indirect effect derived from an hyperphosphorylation of elongation factors such as SPT5 (Cortazar *et al*. 2019).

## DISCUSSION

The study of developmental timing in plants has led to mechanistic dissection of chromatin silencing mechanisms at the gene encoding the Arabidopsis floral repressor FLC. Since quantitative variation of *FLC* expression affects the reproductive strategy of Arabidopsis, any molecular variation can be subject to strong evolutionary selection. *FLC* is thus an excellent system to dissect RNA processing, chromatin regulation, and their interconnections and molecular feedbacks that generate low and high transcriptional states that underpin adaptively important variation in transcriptional output.

This work identifies the importance of APRF1, a component of an RNA Pol II termination machinery, in *FLC* regulation. We identify LD, a protein characterized as a flowering regulator nearly 20 years ago as structurally related to yeast Ref2 and metazoan PNUTS proteins. Given its interaction partners, LD thus acts as a bridge between chromatin modifiers and the Pol II-machinery (Figure 6A). We propose that actively transcribed *FLC* chromatin is enriched with H3K4me1, which promotes RNA Pol II processivity, i.e. the likelihood of transcription reaching the end of the gene. Any pause in RNA Pol II functioning, for example coincident with the formation of the R-loop during *COOLAIR* transcription, would stimulate the 3’ processing machinery (carried along in the RNA Pol II supercomplex) to polyadenylate the transcript at the proximal site. PAS recognition would signal activation of the APRF1-phosphatase module and trigger a conformational change on RNA Pol II (as proposed by Carminati *et al*. 2023) to activate FLD function (Figure 6B). RNA Pol II downstream of the PAS would proceed slowly due to the conformational change, causing FLD to remove H3K4me1 co-transcriptionally until the RNA Pol II is terminated by 5’-3’ XRNs (Figure 6C). This would create a less processive chromatin environment for subsequent rounds of transcription. Each round of proximal polyadenylation-termination-H3K4me1 removal would reinforce the next round of transcription, creating an intrinsic feedback loop in the mechanism (Menon *et al*. 2023). Therefore, we propose transcription termination events contribute to the definition of chromatin domains around genes preventing future transcriptional readthrough.

**Figure 6.**
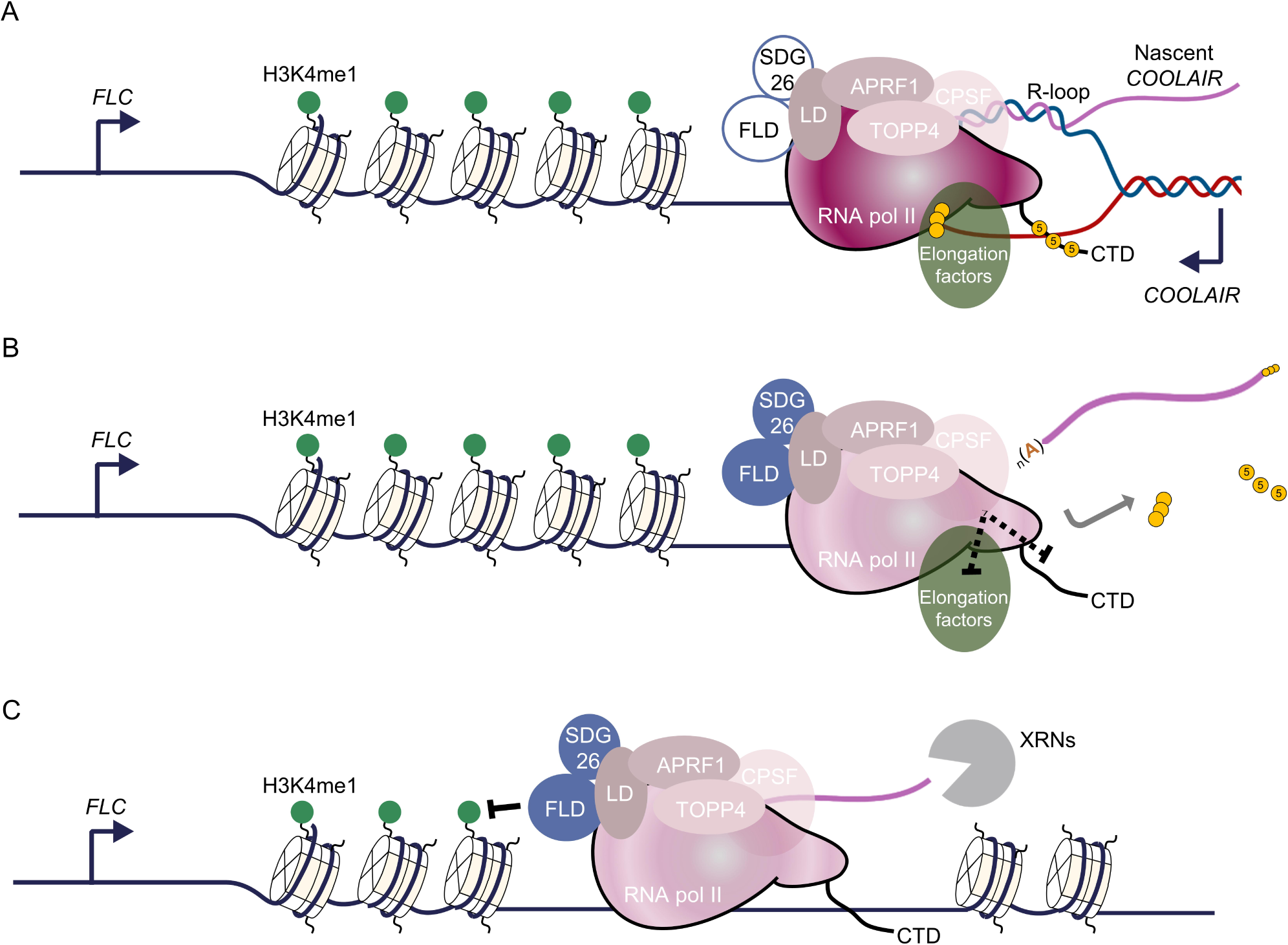
Proposed model for transcription-mediated chromatin silencing. (**A**) Open *FLC* chromatin represented by white nucleosomes and marked with H3K4me1 (green circles) is actively transcribed by RNA Pol II machinery (solid maroon), which carries a non-active FLD complex (blue circle) as well as the 3’-end processing machinery (CPSF), including the phosphatase module formed by APRF1-LD TOPP4, and elongation factors (green oval). Pol II CTD and elongation factors harbour some posttranslational modifications including phosphorylation (yellow circles). Nascent *COOLAIR* forms an R-loop in the 3’ end of the locus (Sun *et al*. 2013, Xu *et al*. 2021b). (**B**) Formation of the R-loop stimulates the 3’-end processing machinery to terminate transcription at the proximal PAS. This termination is also signalled by the phosphatase module to Pol II via dephosphorylation of either elongating factors or the CTD or both (dashed line). PAS recognition triggers a conformational change on RNA Pol II, illustrated by a solid-to-pale maroon colour change, which also activates FLD (now solid blue circle). (**C**) After *COOLAIR* is released, RNA Pol II continues transcribing an uncapped transcript that is the substrate of 5’-3’ exonucleases (XRNs). During this non-productive transcription, the FLD complex co-transcriptionally removes H3K4me1 marks from nucleosomes, creating a less processive chromatin environment for subsequent rounds of transcription.

Intriguingly, our chRNA data detected very clear readthrough effects on the antisense (*COOLAIR*) strand, but also in certain parts of the sense (*FLC*) transcription unit when *APRF1* was disrupted. We thus envisage this mechanism would operate on transcripts from both strands of *FLC*, to first establish and then maintain the transcriptionally silenced state, with the conserved APRF1-LD machinery central to that chromatin silencing mechanism. Proximal termination of *FLC* sense transcription during early embryo development is associated with establishment of the silenced state (Schon *et al*. 2021). Proximally polyadenylated sense transcripts do not accumulate in seedling tissue, but it is possible that these are particularly sensitive to RNA degradation pathways.

A question that arises is whether the CPF phosphatase module functioning at *FLC* associates with the majority of transcribing RNA Pol II, or whether it provides a specialised function on *FLC*. Clearly, some parts of the genome are differentially enriched in co-transcriptional regulators. For example, Arabidopsis genome-wide data indicates that FLD shows a clear enrichment at sites of convergent transcription (Inagaki *et al*. 2021). CPF factors have also been shown to resolve DNA transcription/replication conflicts in plants and humans (Landsverk *et al*. 2020, Baxter *et al*. 2021). Multiple protein phosphatase complexes have been described to promote transcription termination in eukaryotes. The Integrator-PP2C complex comprises 14 subunits (INT1-to-14) with limited conservation in plants (Kirstein *et al*. 2021) participates in both coding and non-coding termination (Hu *et al*. 2023, Wagner *et al*. 2023). Only five of the INT subunits have clear Arabidopsis homologues (INT3/4/7/9/11), with a role in snRNA processing (Liu *et al*. 2016), but none were detected as an APRF1 interactor. The phosphatase module of the CP(S)F/APT and more recently the Restrictor complex have been reported to play a role on transcriptional termination of mRNAs and non-coding RNAs, respectively (Estell *et al*. 2021, Estell *et al*. 2023, Rouvière *et al*. 2023). Homologs of APRF1 (Swd2 and WDR82) are constitutive members of the CPF/CPSF and Restrictor complexes (Lee *et al*. 2010, Schreieck *et al*. 2014, Casanal *et al*. 2017, Estell *et al*. 2021, Estell *et al*. 2023, Rouvière *et al*. 2023, Russo *et al*. 2023), but despite several Zinc-Finger proteins being among the APRF1 interactors none appear to be equivalents of ZC3H4 as part of a plant Restrictor complex (Table S1). Yeast CPF/APT phosphatase modules have other constitutive components such as Pta1, Pti1, and Ssu72 with known orthologs in Arabidopsis [ESP4, Herr *et al*. (2006), Cstf64; Liu *et al*. (2010), Ssu72; Tian *et al*. (2019)] but again these were not found as APRF1 or FLD complex interactors. Thus, we favour from the data we present that APRF1 functions like Swd2 (yeast)/ WDR82 (metazoan) in a CPF-like phosphatase module. Interestingly, in all species studied, the same components are involved in 3’ end processing and transcription termination but they have different affinities. In yeast, the CPF complex contains all enzymatic activities required for mRNA 3’-end processing and transcription termination (cleavage, polyadenylation, dephosphorylation). In humans, the phosphatases are present within the activated 3’-end processing machinery, pulled down on an RNA substrate (Shi *et al*. 2009) but are not constitutive components of CPSF. It remains unclear whether the phosphatase module in plants is constitutively associated with the cleavage and polyadenylation machinery but our mass spectrometry data suggest that it may be a regulated interaction, similar to the human situation (Boreikaite and Passmore 2023). The robust interaction between APRF1 and the CTD phosphatase CPL3 which has been shown to be able to dephosphorylate Ser2, Ser5, and Ser7 (Li *et al*. 2014) may reflect either a simplified termination machinery in plants or the existence of specialized sub-complexes working in an environmentally/developmentally regulated manner.

Another interesting evolutionary difference is the clear sub-functionalization of the Arabidopsis Swd2 homologues *APRF1* and *S2LB* (Birchler and Yang 2022). In yeast, Swd2 is a single copy gene whose product is found in two protein complexes with antagonistic functions: COMPASS (Bae *et al*. 2020) and the mRNA 3’-end processing machinery (Soares and Buratowski 2012, Casanal *et al*. 2017). Arabidopsis orthologs do not play overlapping roles in these two complexes. S2LB interacts *in vivo* with the methyltransferase SDG2 and the structural component of COMPASS WDR5, controlling H3K4me3 levels genome-wide at thousands of gene promoters (Fiorucci *et al*. 2019). In contrast, APRF1 interacts with LD, other components of the FLD complex, and TOPP4. This functional divergence is confirmed through analysis of mutants affecting *FLC* expression. Like *atx1-2* and other mutants in the Arabidopsis COMPASS machinery (Jiang *et al*. 2011), *s2lb* mutants reduce *FLC* and *COOLAIR* expression due to their impaired ability to deposit H3K4me3 at the locus.

The mechanism we describe links transcription termination to histone demethylase activity resulting in graded repression of subsequent transcription. Our accompanying paper then takes this mechanism and describes how it promotes the switch to Polycomb silencing (Menon *et al*. 2023). Only by combining extensive genetics and proteomic analysis described in this paper, with the modelling/experimental validation work described in the accompanying paper, could we fully describe the whole mechanism. A low transcriptional state influenced by proximal termination and H3K4 demethylation reduces the antagonism to Polycomb silencing and leads to a stable PRC2 epigenetic switch, with sufficient feedbacks to maintain the silenced state through DNA replication and cell division. We have generated an animation (weblink) to help explain these molecular feedbacks, and how this mechanism leads to the stable expression of *FLC* in one of two stable expression states. These transcription termination/chromatin silencing mechanisms have proven difficult to dissect at the molecular level in many systems but are likely to be at the basis of many epigenetic switches. There are parallels with the mechanism described here and that of the CPF-triggered heterochromatin silencing in *S. pombe* (Vo *et al*. 2019). 3’ processing has been extensively linked to chromatin silencing in *S. pombe* (Kowalik *et al*. 2015, Grewal 2023), and found to be important for plants to cope with heat shock (Kim *et al*. 2023). In *S. cerevisiae*, a direct connection between a lysine demethylase KDM5 and the CPF was reported (Blair *et al*. 2016). In human cells a clear genome-wide correlation has been found between human FLD and APRF1 homologs (LSD1 and WDR82) and Pol II (Kim *et al*. 2022, Estell *et al*. 2023), and there is a direct relationship between LSD1, and an RNA helicase involved in R-loop resolution and PRC2-mediated silencing (Pinter *et al*. 2021). Continued mechanistic dissection is therefore likely to elaborate generally important concepts in chromatin silencing.

## MATERIALS AND METHODS

### Plant material and growth conditions

All the plants were homozygous for the indicated genotype. The *aprf1-9* mutant seeds (WiscDsLox489-492K11, N858279) were obtained from NASC (Nottingham Arabidopsis Stock Centre). The *s2lb* mutant and the *aprf1-9 s2lb* double mutant, previously described in (Fiorucci *et al*. 2019), were kindly provided by Fredy Barneche. The *atx1-2* was previously described in (Pien *et al*. 2008). Transgenics *fld-4*; *FLD_pro_:3xFLAG-FLD* and *APRF1_pro_:APRF1-3xFLAG* were previously described in (Inagaki *et al*. 2021) and (Qi *et al*. 2022) shared by Soichi Inagaki and Xin-Jian He, respectively.

The *aprf1-10* mutant harbours a 5-nt deletion in *APRF1*. The *aprf1-10* deletion creates a new target for the M*ae*II (target ACGT) restriction enzyme. To generate the *aprf1-10* we employed the CRISPR/Cas9 plasmid pKI1.1R following the protocol described (Tsutsui and Higashiyama 2017). Briefly, pKI1.1R plasmid (Addgene #85808) was linearized by incubating 1.5 µg of the plasmid with A*ar*I restriction enzyme for 16 h, and then dephosphorylated using the alkaline phosphatase rAPid (Roche). A target-specific gRNA was designed using CRIPR-P 2.0 (http://crispr.hzau.edu.cn/CRISPR2). Oligonucleotides harbouring the gRNA target (sgRNA_APRF1_F and sgRNA_APRF1_R; Table S4) were hybridised by slow cooling down from 95-25°C and then phosphorylated using the T4 Polynucleotide Kinase (NEB). The digested plasmid and the hybridised oligonucleotides were ligated using the T4 ligase (NEB) and then transformed in *Escherichia coli* HST08 competent cells (Takara). The sequence integrity of inserts carried by transformants were verified by Sanger sequencing. The plasmid was then transfer to *Agrobacterium tumefaciens* C58C1 strain by electroporation. T_1_ plants carrying the construct were selected on MS media supplemented with 15 µg/ml of Hygromycin. Next generation plants were counter-selected to find transgene-free individuals carrying the homozygous mutation.

Seeds were surface sterilized in 40 % v/v commercial bleach for 10 min and rinsed 4 times with sterile distilled water. Seeds were then sown on standard half-strength Murashige and Skoog (MS) medium (0.22% MS, 8% plant agar) media plates and kept at 4°C in darkness for 3 days before being transferred to long day photoperiod conditions (16 h of light, 8 h dark). All RNA and protein experiments were done using 14-days old seedlings unless otherwise specified.

### Gene expression analyses

Seedlings were harvested, and RNA was extracted with the hot phenol method as previously described (Box *et al*. 2011, Zhu *et al*. 2021). TURBO DNase (Ambion) was used to remove genomic DNA contamination before reverse transcription. cDNA was synthesized using SuperScript IV (Invitrogen) and gene-specific primers (Table S4). qPCR analyses were performed, and data was normalized to the indicated housekeeping gene or genes.

### ChIP

2.5 gr of seedlings were crosslinked with 1% formaldehyde in 1X PBS for 12 min by vacuum infiltration, followed by addition of glycine (final concentration 125 mM) with another 7 min of vacuum infiltration. Tissue was then ground to fine powder with liquid nitrogen. Ground tissue was resuspended in 35 mL of Honda Buffer (20 mM Hepes, 0.44 M sucrose, 1.25 % Ficoll, 2.5% Dextran, 10 mM MgCl_2_, 0.5% Triton X-100, 5 mM DTT, 1x Roche protease inhibitor mixture), filtered through two layers of Miracloth, and centrifuged at 2500 xg for 15 min. Nuclei pellet was then washed once more with 1.6 mL of Honda Buffer.

For histone ChIP, nuclear pellets were resuspended in Nuclei Lysis Buffer (50 mM Tris-HCl pH 8, 10 mM EDTA, 1 % SDS), and sonicated 4 x 5 min (30 sec ON/ 30 sec OFF) using a Diagenode Bioruptor on Medium setting. IP was performed by incubating 140 µl of sonicated chromatin diluted ten times with ChIP dilution buffer (16.7 mM Tris-HCl pH 8, 1.2 mM EDTA, 1.1 % Triton X-100, 167 mM NaCl, 1X cOmplete protease inhibitors) with 15 µl of Protein A-coated Dynabeads (Invitrogen) previously incubated for 2 h with either 2.5 µg of anti-H3 (ab176842), anti-H3K27me3 (ab192985), anti-H3K36me3 (ab9050), or H3K4me1 (ab8895) and incubated overnight at 4°C on a rotator wheel.

For FLAG-FLD and GFP-LD ChIP, nuclei were obtained as described above, but for FLAG-FLD the crosslinking buffer was supplemented with 1.5 mM of EGS [ethylene glycol bis(succinimidyl succinate); ThermoFisher]. Nuclear pellets were suspended in RIPA buffer (50 mM Tris-HCl, 150 mM NaCl, 1% Nonidet P-40, 0.5% NaDeoxycholate, 0.1% SDS, 1x Roche protease inhibitor mixture) and sonicated 5 times x 5 min (30 s ON/ 30 s OFF) with the Bioruptor in high setting. Undiluted chromatin was incubated overnight at 4°C with either 1.5 mg Dynabeads M-270 Epoxy preincubated with 1.5 µl anti-FLAG (Anti-FLAG® M2 / F1804, Merck) for FLAG-FLD or 15 µl Protein-A coated Dynabeads with 2.5 µl anti-GFP (ab290, Abcam), for GFP-LD.

Beads were then washed twice with Low Salt Wash Buffer (150 mM NaCl, 0.1 % SDS, 1% Triton X-100, 2 mM EDTA, 20 mM Tris-HCl pH 8, 1x Roche protease inhibitor mixture), twice with High Salt Wash Buffer (500 mM NaCl, 0.1 % SDS, 1 % Triton X-100, 2 mM EDTA, 20 mM Tris-HCl 8, 1x Roche protease inhibitor mixture), and twice with TE wash buffer (10 mM Tris-HCl pH 8, 1 mM EDTA, 1x Roche protease inhibitor mixture).

For RNA Pol II ChIP, nuclei were obtained as described for histone ChIP, complementing the Honda Buffer with 1x of PhosSTOP (Merck). Nuclear pellet was suspended in 1 mL of TAP buffer (100 mM NaCl, 20 mM Tris-HCl pH 8, 2.5 mM EDTA, 10 % glycerol, 1 % Triton, 1x of PhosSTOP, 1x Roche protease inhibitor mixture) and given 20 strokes with the Dounce Homogenizer. The resulting solution was sonicated 4 x 10 min (15 s ON/ 45 s OFF) with the Bioruptor in low setting. 250 µl of undiluted chromatin was incubated overnight at 4°C with Dynabeads M-270 Epoxy preincubated with anti-Tyr1P (MABE350, Merck), anti-Ser2P (C15200005-50, Diagenode), or anti-Ser5P (C15200007-50, Diagenode). Then beads were washed twice for 15 min with Low Salt and High Salt buffers like for histones but including 1x of PhosSTOP. Then beads were washed for 15 min with the LiCl buffer (250 mM LiCl, 0.5 % NP40, 2.5 mM EDTA, 0.05 % NaDeoxycholate, 20 mM Tris-HCl pH 8, 1x of PhosSTOP, 1x Roche protease inhibitor mixture), and the TE buffer (same as for histones plus 1x PhosSTOP). In all cases, after IP, DNA was then eluted and reverse-crosslinked by incubating the beads at 95°C for 10 min in presence of 100 µl of 10 % Chelex resin (BioRad), treated with Proteinase K (Roche) for 1 h at 45°C, and incubated again at 95°C for 10 min to inactivate the Proteinase K. Finally, DNA was purified using the ChIP DNA Clean & Concentrator kit (Zymo Research).

### CrossLinked Nuclear ImmunoPrecipitation and Mass Spectrometry (CLNIP-MS)

10-days-old *APRF1_pro_:APRF1-3xFLAG* (Qi *et al*. 2022) and Col-0 (control) seedlings were crosslinked with 1% formaldehyde in 1X PBS for 10 min. Three biological replicates for each genotype were used. 2 g of tissue per biological replicate was ground to a fine powder and resuspend in 30 mL of Honda Buffer, supplemented with 1 mM phenylmethylsulfonyl fluoride (PMSF). The suspension was filtered through a double layer of Miracloth and centrifuged at 2,000 g for 15 min at 4°C. The nuclei pellet was washed once in 5 mL of Honda buffer then purified on a Percoll density gradient as follows: 2 mL of 75% Percoll (Merck, P7828) in Honda buffer topped with 2 mL of 40% Percoll in Honda buffer topped with the nuclei pellet resuspended in Honda buffer in a 15 mL tube. Purified nuclei were obtained in between the layers containing 40% and 75% Percoll after centrifuging at 7,000 g for 30 min at 4°C and washed once more in 6 mL Honda buffer. The nuclei pellet was resuspended in 350 μL of Benzonase buffer (50 mM Tris pH 8.0, 1 mM MgCl2, and 1X cOmplete protease inhibitors), and incubated with 1 μL of Benzonase (Millipore, 70746) for 40 min at 4°C. Nuclei were then incubated for 30 min at 4°C after adding 1% SDS, then diluted with ChIP dilution buffer to a concentration of 0.5% SDS in the samples, then sonicated using the Bioruptor in Medium setting for 3 cycles of 5 min (30 sec on/ 30 sec off). Samples were centrifuged at 10,000 g for 1 min, and the supernatant was diluted with ChIP dilution buffer to a concentration of 0.1% SDS in the sample. IP was performed overnight at 4°C after adding the antibody-beads complex. 1.5 µg of anti-FLAG (Sigma, F1804) antibody was coupled to 1.5 mg of M-270 epoxy Dynabeads (Invitrogen, 14311D) following the manufacturer’s procedure and used per IP reaction. After IP, samples were washed with 1 mL of IP wash buffer (50 mM Tris pH 8.0, 150 mM NaCl, 1% Triton XL100, and 0.5% IGEPAL CA-630) 4 times for 5 min each, and then resuspend in 50 μL of SDS buffer (20 mM Tris pH 8.0 and 2% SDS), and heated to 90°C for 15 min. Then, the samples were separated from the beads and proteins precipitated based on (Pankow *et al*. 2016) by adding 1.1/4.4 chloroform/methanol mix. The protein pellet was then washed twice with methanol, once with acetone, and air dried.

For Mass Spectrometry, protein pellets were resuspended in 50 µl of 1.5% sodium deoxycholate (SDC; Merck) in 0.2 M EPPS-buffer (Merck), pH 8.5 and vortexed under heating. Cysteine residues were reduced with dithiothreitol, alkylated with iodoacetamide, and the proteins digested with trypsin in the SDC buffer according to standard procedures. After the digest, the SDC was precipitated by adjusting to 0.2% trifluoroacetic acid (TFA), and the clear supernatant subjected to C18 SPE using home-made stage tips with C18 Reprosil_pur 120, 5 µm. Aliquots were analysed by nanoLC-MS/MS on an Orbitrap Eclipse™ Tribrid™ mass spectrometer coupled to an UltiMate® 3000 RSLCnano LC system (Thermo Fisher Scientific, Hemel Hempstead, UK). The samples were loaded onto a trap cartridge (PepMap™ Neo Trap Cartridge, C18, 5um, 0.3×5mm, Thermo) with 0.1% TFA at 15 µl min-1 for 3 min. The trap column was then switched in-line with the analytical column (Aurora Frontier TS, 60 cm nanoflow UHPLC column, ID 75 µm, reversed phase C18, 1.7 µm, 120 Å; IonOpticks, Fitzroy, Australia) for separation at 55°C using the following gradient of solvents A (water, 0.1% formic acid) and B (80% acetonitrile, 0.1% formic acid) at a flow rate of 0.26 µl min-1: 0-3 min 1% B (parallel to trapping); 3-10 min increase B (curve 4) to 8%; 10-102 min linear increase B to 48; followed by a ramp to 99% B and re-equilibration to 0% B, for a total of 140 min runtime. Mass spectrometry data were acquired with the FAIMS device set to three compensation voltages (−35V, −50V, −65V) at standard resolution for 1.0 s each with the following MS settings in positive ion mode: OT resolution 120 K, profile mode, mass range m/z 300-1600, normalized AGC target 100%, max inject time 50 ms; MS2 in IT Turbo mode: quadrupole isolation window 1 Da, charge states 2-5, threshold 1e4, HCD CE = 30, AGC target standard, max. injection time dynamic, dynamic exclusion 1 count for 15 s with mass tolerance of ±10 ppm, one charge state per precursor only.

The mass spectrometry raw data were processed and quantified in Proteome Discoverer 3.1 (ThermoFisher) using the search engine CHIMERYS (MSAID, Munich, Germany); all mentioned tools of the following workflow are nodes of the proprietary Proteome Discoverer (PD) software. The Arabidopsis TAIR10 protein database (arabidopsis.org; 32785 entries) was imported into PD adding a reversed sequence database for decoy searches; in the same way, a small customs database with the APRF1-FLAG protein sequence and a database for common contaminants (maxquant.org, 245 entries) was also included. The CHIMERYS database search was performed with the inferys_3.0.0_fragmentation prediction model, a fragment tolerance of 0.3 Da, enzyme trypsin with 2 missed cleavages, variable modification oxidation (M), fixed modification carbamidomethyl (C) and FDR targets 0.01 (strict) and 0.05 (relaxed). The workflow included the Minora Feature Detector with min. trace length 5, S/N 2.5, PSM confidence high. The consensus workflow in the PD software was used to evaluate the peptide identifications and to measure the abundances of the peptides based on the LC-peak intensities. For identification, an FDR of 0.01 was used as strict threshold, and 0.05 as relaxed threshold.

For quantification, three replicates of *APRF1_pro_:APRF1-FLAG* and Col-0 were measured. In PD3.1, the following parameters were used for ratio calculation: normalisation on total peptide abundances, protein abundance-based ratio calculation using the top three most abundant peptides, missing values imputation by low abundance resampling, hypothesis testing by t-test (background based), adjusted p-value calculation by BH-method. The results were exported into a Microsoft Excel table including data for protein abundances, ratios, p-values, number of peptides, protein coverage, the CHIMERYS identification score and other important values. The mass spectrometry proteomics data have been deposited to the ProteomeXchange Consortium via the PRIDE (Perez-Riverol *et al*. 2022) partner repository with the dataset identifier PXD049114 and 10.6019/PXD049114.

### Protein co-immunoprecipitation in *Nicotiana benthamiana*

To generate *APRF1_pro_:APRF1-mVENUS* and *TOPP4_pro_:TOPP4-3xFLAG* the genomic region of both genes including a region of around 1.5 kb upstream their transcription start sites were amplified by PCR and cloned by the InFusion cloning (Takara) into the pCAMBIA1300 in frame with the coding sequences of mVENUS or the 3xFLAG peptide, respectively, using the primers listed in Table S4. After verification by sequencing, both transgenes were transferred to the strain GV3101 of *Agrobacterium tumefaciens.* As a negative control, we used FLAG-tagged version of the Arabidopsis TCP14, a transcription factor involved in plant immunity (Wessling *et al*. 2014), kindly shared by Jonathan D. G. Jones (The Sainsbury Laboratory). Before co-infiltration, protein levels of individual proteins were verified by single agroinfiltrations. Overnight grown bacteria were used to agroinfiltrate adjusting the OD600 to their protein levels. Cells were resuspended in the Infiltration Buffer (10 mM MES pH 5.6, 10 mM MgCl2, 1 mM acetosyringone) and used to agroinfiltrated *N. benthamiana* leaves. Co-infiltrated leaves with either APRF1-mVENUS + TOPP4-3xFLAG or APRF1-mVENUS + TCP14-FLAG were grown 2 more days before collecting the material. Around 0.85 gr of infiltrated tissue was harvested and ground to a fine powder with liquid nitrogen and homogenized in ice-cold extraction buffer (10 % glycerol, 25 mM Tris-HCl pH 7.5, 1 mM EDTA, 150 mM NaCl, 2 %PVP, and 0.2 % Tween-20). The lysate was homogenized, washed twice, and filtered through Miracloth. The supernatant was incubated with 30 μL of washed anti-FLAG M2 Affinity Gel (A2220, Millipore) for 2 hours at 4°C in rotation. The beds were washed four times with IP wash buffer (25 mM Tris-HCl pH 7.5, 1 mM EDTA, 150 mM NaCl, 0.2 % Tween-20, 1 mM DTT, 1x cOomplete protease inhibitors) at 4°C, and resuspended in SDS-loading buffer with 10 mM DTT. Proteins were released and denatured after incubation at 95°C for 7 min and resolved by SDS-PAGE. Anti-FLAG M2-HRP (A8592, Sigma), or anti-GFP (sC9996HRP, Santa Cruz) antibodies were used for Western blot to detect TOPP4-FLAG (expected size: 39 kDa), TCP14-FLAG (52 kDa), or APRF1-mVENUS (62 kDa), respectively. The chemiluminescence substrate SuperSignal West Pico (34580) was used for FLAG immunoblots and SuperSignal West Femto (34094, Thermo) for GFP blots. Uncropped blots are shown in Figure S8.

### Preparation of Chromatin-bound RNA

Chromatin-bound RNA was isolated as previously described (Wu *et al*. 2016). Nuclei from 2-2.5 gr of non-crosslinked seedlings were obtained with Honda Buffer, supplemented with 20 U/mL RNase inhibitor RNase Out (Invitrogen), 1 mM PMSF, and 50 ng/µL of yeast tRNA. Nuclear pellet was rinsed with 500 µL of resuspension buffer (20 mM Tris pH 8, 75 mM NaCl, 0.5 mM EDTA, 1 mM DTT, 0.125 mM PMSF, 50 % glycerol, 1x Roche complete, 20 U/mL RNase Out) and centrifuged at 4000 xg at 4°C for 3 min. Nuclei pellet was weighed and resuspended in an equal volume of Resuspension Buffer. The suspension was then washed with two volumes of Urea Wash Buffer (20 mM Tris pH8, 300 mM NaCl, 7.5 mM MgCl_2_, 0.25 mM EDTA, 1 mM DTT, 1 M Urea, 1 % NP-40, 1x Roche complete, 20 U/mL RNase Out), pipetting up and down 30 times, and spun at 8,000 xg for 1 min at 4°C. Nuclei pellet was resuspended again with 1 volume of Resuspension buffer and washed with 1 volume of Urea Wash Buffer, pipetting 30 times up and down, and spun for 1 min at 8,000 xg and 4°C. Finally, nuclear pellet was dissolved in 1 mL of TRIzol (Invitrogen), adding 0.2 mL of chloroform and shaking vigorously by hand for 15 s. Then the suspension was incubated at RT for 2 min and centrifuged at 12,000 xg for 15 min at 4°C. The aqueous phase was taken and mixed with an equivalent volume of Phenol:Chloroform:Isoamyl alcohol (25:24:1, Sigma), shaken for 10 min at room temperature, and centrifuged at RT and 12,000 xg for 10 min. The aqueous phase was then transferred to another tube and followed two precipitations, first with isopropanol, sodium acetate, and RNA GlycoBlue and another one with LiCl. Finally, RNA was dissolved and treated with DNase Turbo (Ambion) and used as template for reverse transcription with gene-specific primers (Table S4).

### Quant-seq

For Quant-seq experiments, total RNA was isolated as for qPCR and further cleaned up using the Qiagen RNeasy miniprep kit (74106). Library preparation, sequencing, and data analysis were carried out by Lexogen GmbH (Austria). Sequencing-ready libraries were generated from 100 ng of input RNA using a QuantSeq 3’ mRNA-Seq Library Prep Kit REV for Illumina (015UG009V0271) following standard procedures. RNA integrity, and Indexed libraries quality were assessed on a Fragment Analyzer device (Agilent Technologies) using a DNF-471 RNA Kit and HS-DNA assay, respectively. Libraries were quantified using a Qubit dsDNA HS assay (Thermo Fisher). A sequencing-ready pool of indexed libraries were sequenced on an Illumina NextSeq 2000 with a 100-cycle cartridge using the Custom Sequencing Primer (CSP). FastQC version v0.11.7 was used to verify the read quality and cutadapt version 1.18 (Martin 2011) for read adapter trimming. Clean reads were mapped to the latest version of the Arabidopsis genome (TAIR10) with a splice-aware aligner STAR version 2.6.1a (Dobin *et al*. 2013). Differentially expressed genes (DEGs) between Col*FRI* and *aprf1-9 FRI* and Col-0 and *fld-4* are listed in Tables S2 and S3, respectively.

For enrichment of *FLC* and selected control genes, 4,861 synthetic 80-nt biotinylated RNA probes were synthesized, complementary to 32 padded gene sequences at 2x bp tiling density (±1kb padding) (*mybaits*; ArborBiosciences; Supplemental Dataset 1). Selected libraries were pooled equimolar and in-solution target capture was carried out using the manufacturers standard sensitivity protocol with a bait annealing temperature of 65°C. The bait enriched library pool was sequenced on an Illumina NextSeq 2000 with a 100-cycle cartridge using the Custom Sequencing Primer (CSP). Raw reads have been deposited on Short Read Archive (SRA) under the references PRJNA978558 and PRJNA1076161.

### plaNET-seq

Nascent transcript isolation was adapted from (Kindgren *et al*. 2020). 3 gr of Arabidopsis seedlings were flash frozen in liquid nitrogen and extracted with NUC1 [0.4 M sucrose, 10 mM Tris–HCl pH 8.0, 10 mM MgCl_2_, 5 mM β-mercaptoethanol, proteinase inhibitor (Complete; Roche), phosphatase inhibitor (PhosSTOP; Roche) and RNase inhibitor (RNasin; Promega)]. Once homogeneous, samples were centrifuged at 5000 g for 20 minutes and the pellet was washed with 1 ml NUC2 buffer (0.25 M sucrose, 10 mM Tris–HCl pH 8.0, 10 mM MgCl_2_, 5 mM β-mercaptoethanol, proteinase inhibitor, phosphatase inhibitor, RNase inhibitor and 0.3 % Tween-20). Nuclei were suspended in 0.3 ml NUC3 buffer (1.7 M sucrose, 10 mM Tris–HCl pH 8.0, 2 mM MgCl_2_, 5 mM β-mercaptoethanol, proteinase inhibitor tablet, phosphatase inhibitor, RNase inhibitor (Recombinant RNasin 20 U/ml; Promega) and 0.15 % Tween-20) and carefully layered over 0.9 ml of NUC3 in prechilled microcentrifuge tubes before centrifugation at 16000 g for 60 min at 4°C. Purified nuclei were lysed in 1.5 ml plaNET-seq lysis buffer [0.3 M NaCl, 20 mM Tris–HCl pH 7.5, 5 mM MgCl_2_, 5 mM DTT, proteinase inhibitor, phosphatase inhibitor, RNase inhibitor, 0.5% Tween-20 and DNaseI (400 U/ml; Roche)] at 4°C shaking at 2000 rpm. Lysate was centrifuged at 10000 g at 4°C for 10 minutes and supernatant was transferred to Dynabeads M-270 (Invitrogen) coupled to either CTD Ser2P (C15200005; Diagenode) or Ser5P (C15200007; Diagenode) antibodies. After 2 hours incubation at 4°C, immunocomplexes were washed gently six times with wash buffer (0.3 M NaCl, 20 mM Tris-HCl pH 7.5, 5 mM MgCl_2_, 5 mM DTT, proteinase inhibitor RNase inhibitor and phosphatase inhibitor) and dissolved in 1 ml TRIzol (Invitrogen), followed by isolation of the nascent RNA with on column DNA digestion (RNA microprep kit; Direct-zol).

For plaNET-seq library construction, 100 ng of nascent RNA was used as input to construct plaNETseq libraries using NEXTflex Small RNA-seq kit v3 (PerkinElmer) with a modified protocol. After 3’ adapter ligation, RNA was fragmented by incubation with alkaline solution (100 mM NaCO3 pH 9.2, 2 mM EDTA) at 95°C for 5 minutes (Churchman & Weissman 2012), followed by clean up (RNAclean XP beads; Beckman Coulter), PNK treatment (NEB) for 20 min at 37°C and reannealing of the RT-primer (8 μM). Library construction continued from the adapter inactivation step of the manufacturer’s protocol. Libraries were quantified using a Qubit dsDNA HS assay (Thermo Fisher). A sequencing-ready pool of indexed libraries were sequenced on an Illumina Xten PE150 at Beijing Genomics Institute. Raw reads have been deposited on SRA under the reference PRJNA1076151.

For plaNET-seq data analysis, Unique Molecular Identifiers (UMIs) were first trimmed from the read and appended to the read name with UMI-tools v1.1.1 (Smith *et al*. 2017), followed by adapter and read quality trimming with trimmomatic v0.39 (Bolger *et al*. 2014). R2 reads were mapped to the Arabidopsis genome (TAIR10) with a splice-aware aligner STAR version 2.7.10a (Dobin *et al*. 2013). PCR duplicates were filtered from the alignment files with UMI-tools, low mapping quality reads were removed (MAPQ>10 samtools v1.9) and reads were flipped to restore the original RNA read strand orientation. Read 3’ends that overlap with 5’ and 3’ splice sites (and likely represent co-transcriptional splicing intermediates) were removed before generating strand specific coverage files for visualisation of nascent transcripts. For generate the metaplots, gene models from Araport 11 were used to define the TSS and the TTS as described in (Kindgren *et al*. 2020). The gene list was filtered to remove genes overlapping features within 500 bp of the TSS or TTS. For the remaining genes average signal for each position was calculated around the TSS or TTS (±500 bp) and divided into 5 bp bins. For each position 0.01% of extreme values were trimmed before averaging. The mean coverage for each genotype was plot for the given genomic interval with the shaded area indicating the 95% confidence interval for the mean. For the Readthrough analyses, the total binned signal between 150-450 bp downstream of the TTS was calculated for each sample to generate the readthrough signal. The gene 3’end signal was calculated by taking the total signal from a random but equal number of bins within the 3’end 1kb-0.1kb upstream of the TTS. Readthrough rate was expressed as a ratio of readthrough signal relative to 3’ end signal for each replicate. One-way ANOVA was performed to determine statistical significance of the differences between genotypes.

### AlphaFold2 protein interaction prediction

AlphaFold models were predicted using the Colab notebook running a slightly simplified version of AlphaFold v2.3.2. (Jumper *et al*. 2021). For the Arabidopsis LD–APRF1 complex LD aa566-661 and APRF1 aa1-330 were used. For the human PNUTS–WDR82 complex PNUTS aa380-530 and WDR82 aa1-313, and for the yeast Ref2–Swd2 complex Ref2 aa406-533 and Swd2 aa1-329 were used. The number of recycles were set to 3 and models included a final relaxation stage. PAE plots indicating the quality of the prediction and LDDT plots are shown in Figure S15. Final models were imported into and colour figures were prepared with PyMOL (v2.1, Schrödinger).

### Bioinformatic analyses

To design the sgRNA to edit *APRF1* via CRISPR/Cas9 we used the CRISPR-P 2.0 (http://crispr.hzau.edu.cn/cgi-bin/CRISPR2/CRISPR), selecting the canonical NGG PAM motif, and the Arabidopsis TAIR10 as a target genome. Protein alignments were performed with MEGA X (Kumar *et al*. 2018) using the MUSCLE (Edgar 2004) algorithm with default parameters, and shaded with the “Colour Align Conservation” tool from the Sequence Manipulation Suite (Stothard 2000). To find putative PNUTS homologs in Arabidopsis, BLASTP (Altschul *et al*. 1997), PSI-BLAST, and DELTA-BLAST (Boratyn *et al*. 2012) searches were performed at NCBI site using an alignment score threshold of 80 for DELTA-BLAST, and default parameters for the rest. Individual protein structure predictions were retrieved from the Alphafold website (https://alphafold.ebi.ac.uk/). Statistical analyses are detailed in every Figure and were carried out using GraphPad Prism version 9.0.0 for Windows, GraphPad Software.

## ACKNOWLEDGEMENTS

For genetic materials, we are indebted to Fredy Barneche for providing seeds of *s2lb* and *aprf1-9 s2lb* (Fiorucci *et al*. 2019), Soichi Inagaki for sharing their *3xFLAG-FLD* [in both (*fld-4; FRI*) and (*fld-4*; *fri*) backgrounds] transgenic lines (Inagaki *et al*. 2021), and Xin-Jian He for sharing the *APRF1-3xFLAG* line (Qi *et al*. 2022). We are indebted to Jianhua Huang for his precious help and expertise in transient protein co-expression and for providing the TCP14-FLAG control line, to Yusheng Zhao for initiating the introgression of the GFP-LD line into Col*FRI* and to Minglei Yang for writing the code used to analyse Quant-seq data. Authors would also thank Shuqin Chen, Aida Sánchez, and Tina Zhang for their excellent technical assistance. This work was funded by the European Research Council Advanced Grant (EPISWITCH, 833254), Wellcome Trust (210654/Z/18/Z) and the Royal Society Professorship (RP\R1\180002) to Caroline Dean; BBSRC Institute Strategic Programmes (BB/J004588/1 and BB/P013511/1) and the Medical Research Council (MRC) as part of United Kingdom Research and Innovation (MRC file reference number MC_U105192715) and the Wellcome Trust (225217/Z/22/Z) to Lori A Passmore. Eduardo Mateo-Bonmatí would like to thank grant RYC2021-030895-I, funded by MCIN/AEI and by European Union NextGenerationEU/PRTR.

