## Supplemental Figures for "A CPF-like phosphatase module links transcription termination to chromatin silencing"

### Slide 1
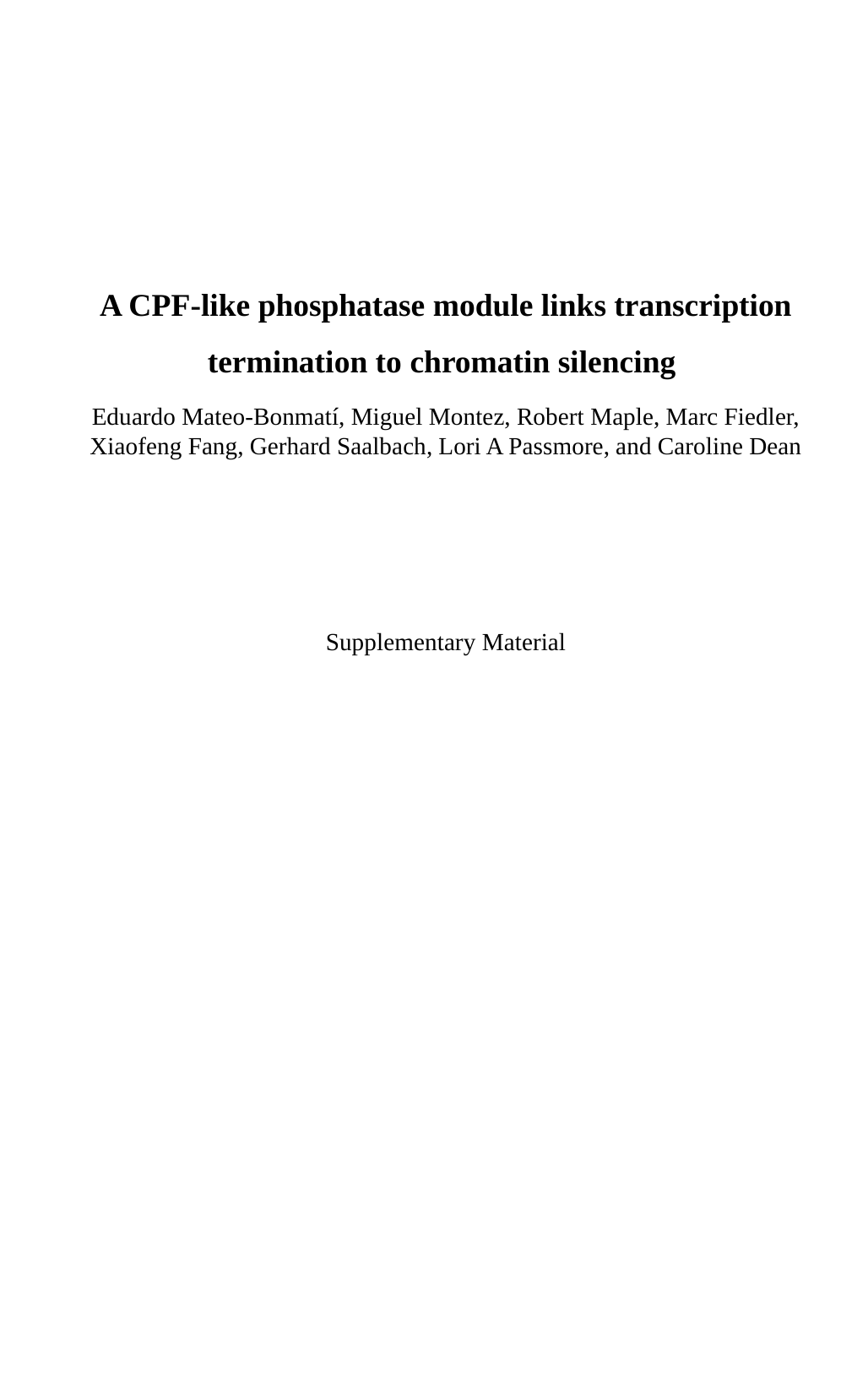

A CPF-like phosphatase module links transcription termination to chromatin silencing
Eduardo Mateo-Bonmatí, Miguel Montez, Robert Maple, Marc Fiedler, Xiaofeng Fang, Gerhard Saalbach, Lori A Passmore, and Caroline Dean
Supplementary Material

### Slide 2
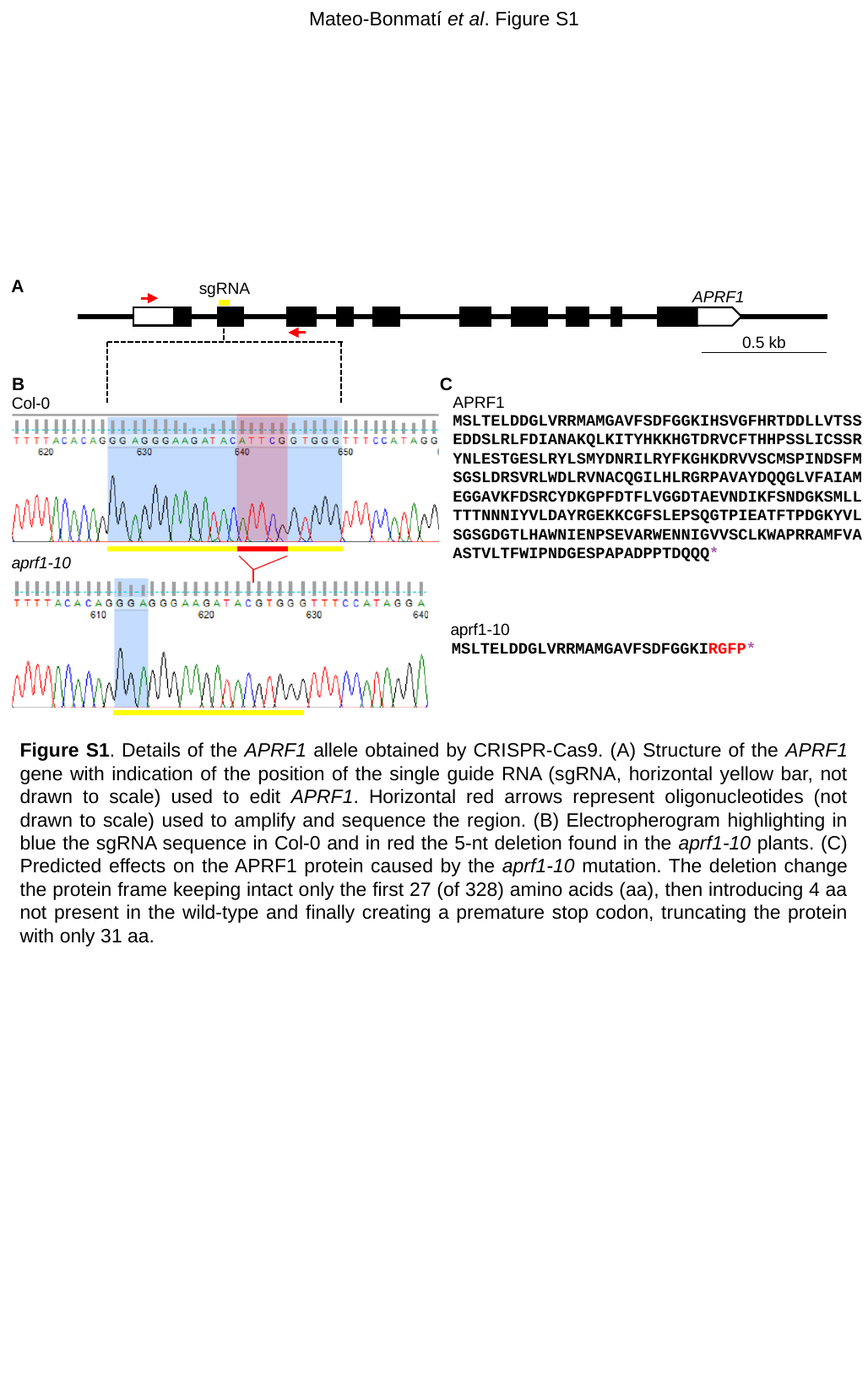

Mateo-Bonmatí et al. Figure S1
A
sgRNA
APRF1
0.5 kb
aprf1-10
B
C
APRF1
Col-0
MSLTELDDGLVRRMAMGAVFSDFGGKIHSVGFHRTDDLLVTSSEDDSLRLFDIANAKQLKITYHKKHGTDRVCFTHHPSSLICSSRYNLESTGESLRYLSMYDNRILRYFKGHKDRVVSCMSPINDSFMSGSLDRSVRLWDLRVNACQGILHLRGRPAVAYDQQGLVFAIAMEGGAVKFDSRCYDKGPFDTFLVGGDTAEVNDIKFSNDGKSMLLTTTNNNIYVLDAYRGEKKCGFSLEPSQGTPIEATFTPDGKYVLSGSGDGTLHAWNIENPSEVARWENNIGVVSCLKWAPRRAMFVAASTVLTFWIPNDGESPAPADPPTDQQQ*
aprf1-10
MSLTELDDGLVRRMAMGAVFSDFGGKIRGFP*
Figure S1. Details of the APRF1 allele obtained by CRISPR-Cas9. (A) Structure of the APRF1 gene with indication of the position of the single guide RNA (sgRNA, horizontal yellow bar, not drawn to scale) used to edit APRF1. Horizontal red arrows represent oligonucleotides (not drawn to scale) used to amplify and sequence the region. (B) Electropherogram highlighting in blue the sgRNA sequence in Col-0 and in red the 5-nt deletion found in the aprf1-10 plants. (C) Predicted effects on the APRF1 protein caused by the aprf1-10 mutation. The deletion change the protein frame keeping intact only the first 27 (of 328) amino acids (aa), then introducing 4 aa not present in the wild-type and finally creating a premature stop codon, truncating the protein with only 31 aa.

### Slide 3
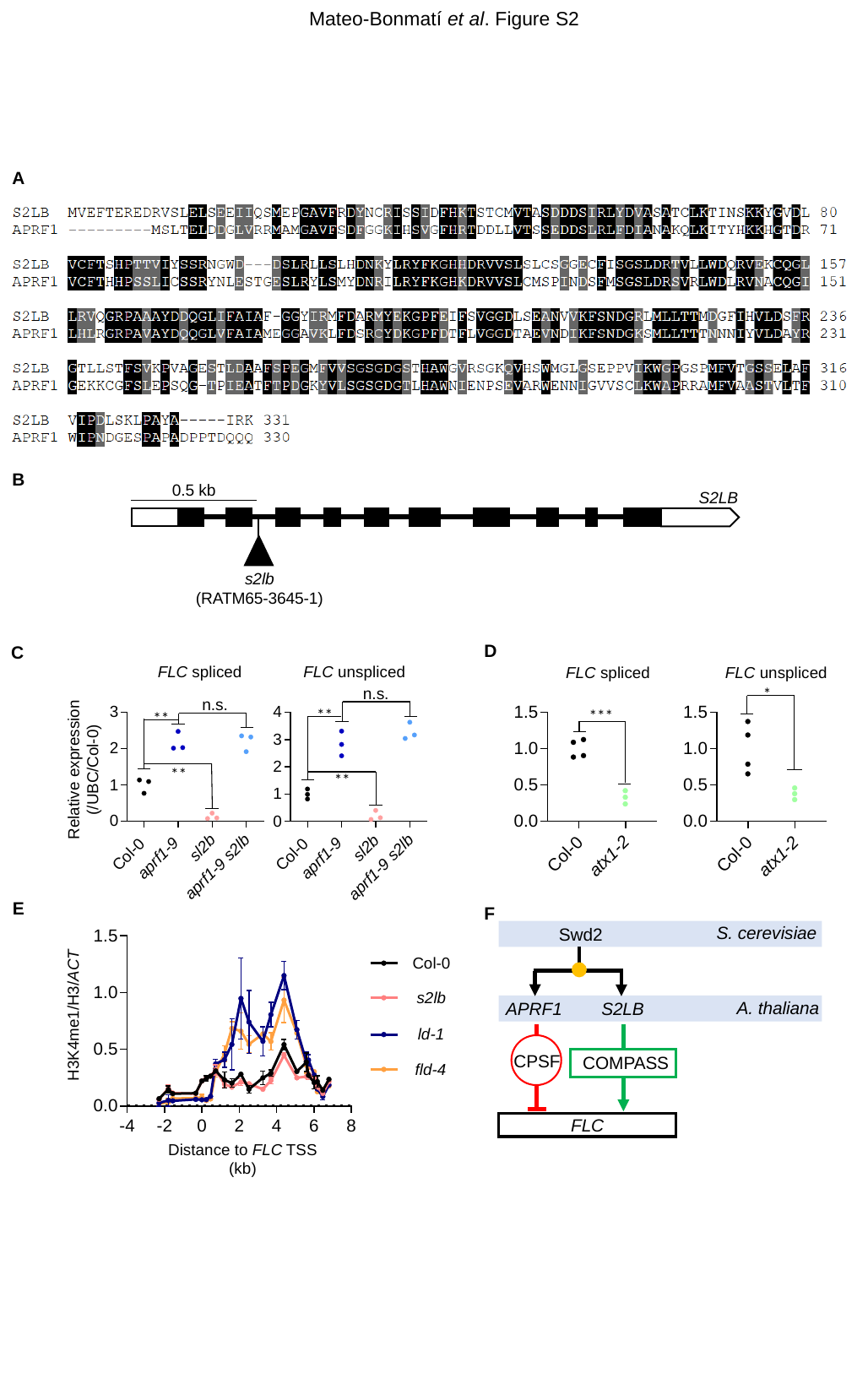

Mateo-Bonmatí et al. Figure S2
A
B
0.5 kb
S2LB
s2lb
(RATM65-3645-1)
D
C
FLC unspliced
FLC spliced
FLC unspliced
FLC spliced
n.s.
*
n.s.
**
***
**
Relative expression (/UBC/Col-0)
**
**
E
F
S. cerevisiae
Swd2
S2LB
APRF1
CPSF
COMPASS
FLC
Col-0
s2lb
A. thaliana
H3K4me1/H3/ACT
ld-1
fld-4
Distance to FLC TSS (kb)

### Slide 4
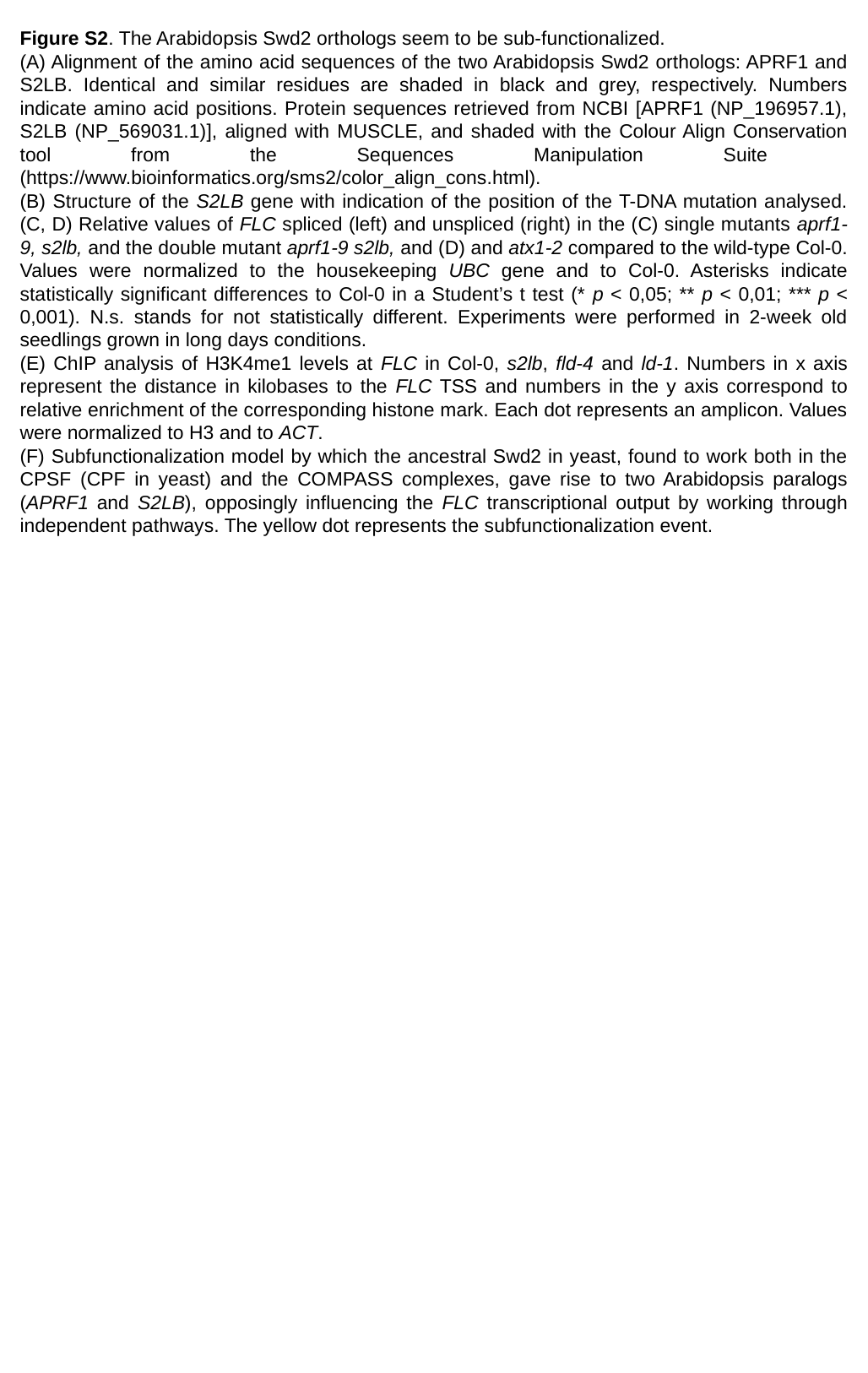

Figure S2. The Arabidopsis Swd2 orthologs seem to be sub-functionalized.
(A) Alignment of the amino acid sequences of the two Arabidopsis Swd2 orthologs: APRF1 and S2LB. Identical and similar residues are shaded in black and grey, respectively. Numbers indicate amino acid positions. Protein sequences retrieved from NCBI [APRF1 (NP_196957.1), S2LB (NP_569031.1)], aligned with MUSCLE, and shaded with the Colour Align Conservation tool from the Sequences Manipulation Suite (https://www.bioinformatics.org/sms2/color_align_cons.html).
(B) Structure of the S2LB gene with indication of the position of the T-DNA mutation analysed. (C, D) Relative values of FLC spliced (left) and unspliced (right) in the (C) single mutants aprf1-9, s2lb, and the double mutant aprf1-9 s2lb, and (D) and atx1-2 compared to the wild-type Col-0. Values were normalized to the housekeeping UBC gene and to Col-0. Asterisks indicate statistically significant differences to Col-0 in a Student’s t test (* p < 0,05; ** p < 0,01; *** p < 0,001). N.s. stands for not statistically different. Experiments were performed in 2-week old seedlings grown in long days conditions.
(E) ChIP analysis of H3K4me1 levels at FLC in Col-0, s2lb, fld-4 and ld-1. Numbers in x axis represent the distance in kilobases to the FLC TSS and numbers in the y axis correspond to relative enrichment of the corresponding histone mark. Each dot represents an amplicon. Values were normalized to H3 and to ACT.
(F) Subfunctionalization model by which the ancestral Swd2 in yeast, found to work both in the CPSF (CPF in yeast) and the COMPASS complexes, gave rise to two Arabidopsis paralogs (APRF1 and S2LB), opposingly influencing the FLC transcriptional output by working through independent pathways. The yellow dot represents the subfunctionalization event.

### Slide 5
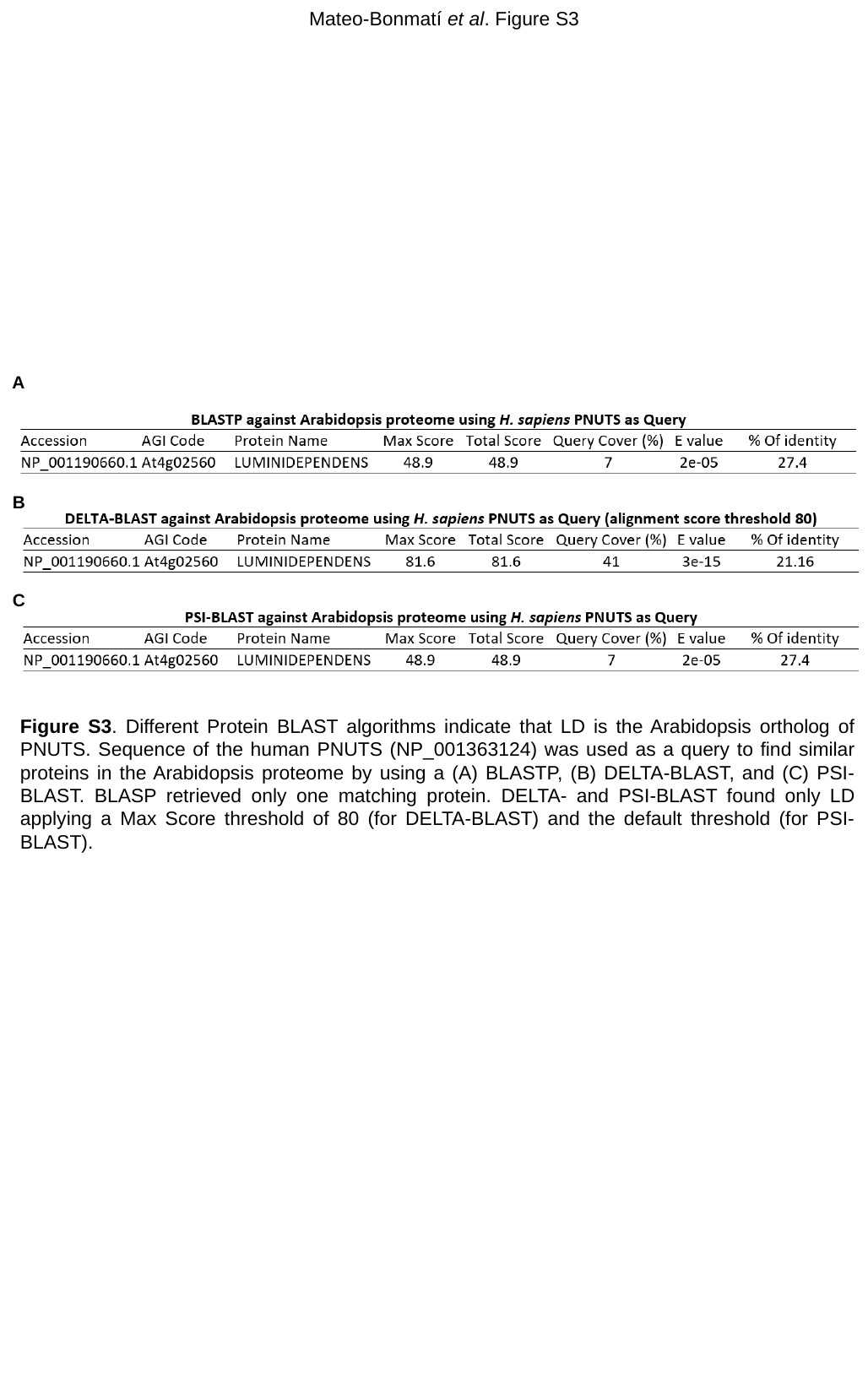

Mateo-Bonmatí et al. Figure S3
A
B
C
Figure S3. Different Protein BLAST algorithms indicate that LD is the Arabidopsis ortholog of PNUTS. Sequence of the human PNUTS (NP_001363124) was used as a query to find similar proteins in the Arabidopsis proteome by using a (A) BLASTP, (B) DELTA-BLAST, and (C) PSI-BLAST. BLASP retrieved only one matching protein. DELTA- and PSI-BLAST found only LD applying a Max Score threshold of 80 (for DELTA-BLAST) and the default threshold (for PSI-BLAST).

### Slide 6
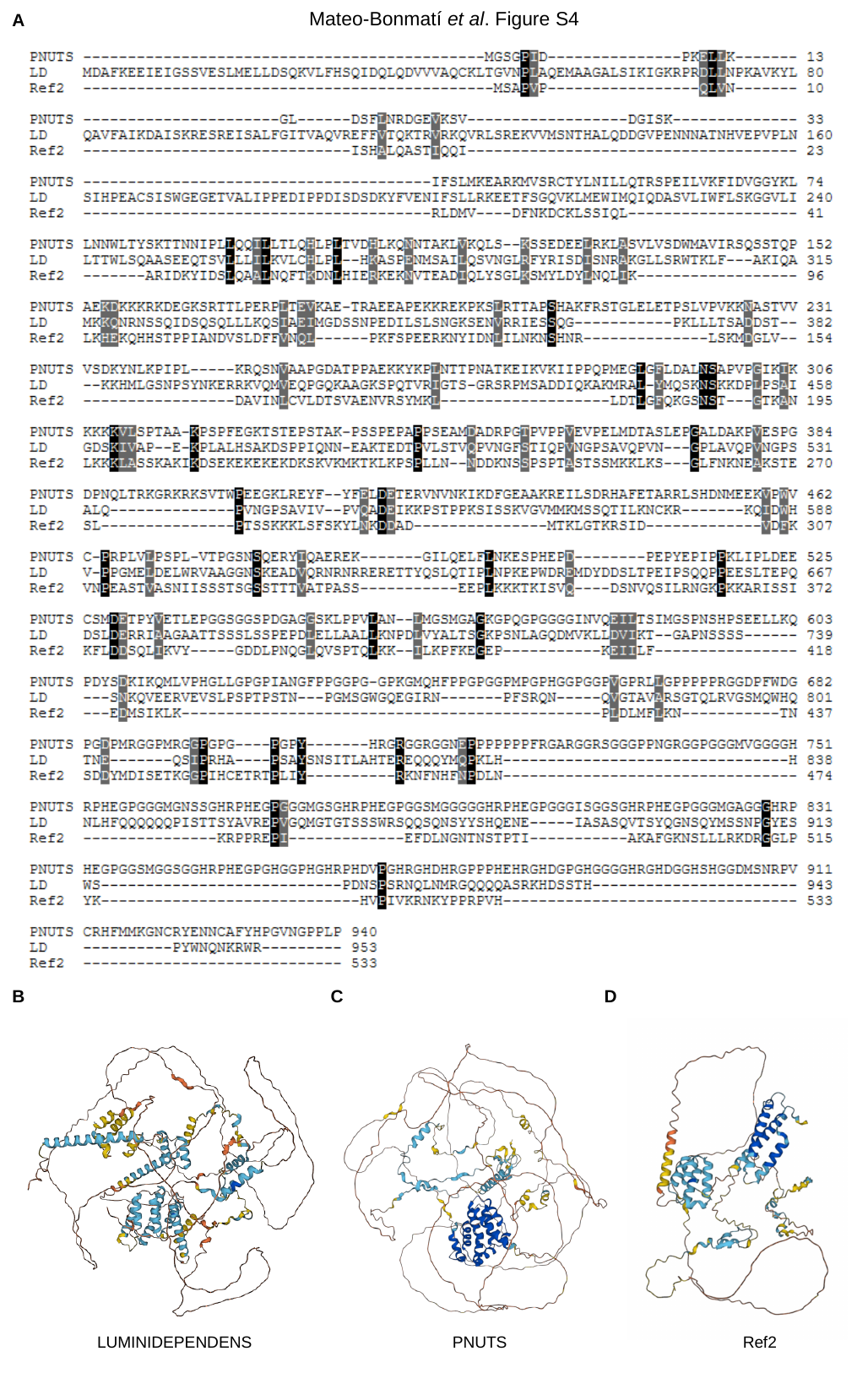

Mateo-Bonmatí et al. Figure S4
A
D
C
B
LUMINIDEPENDENS
PNUTS
Ref2

### Slide 7
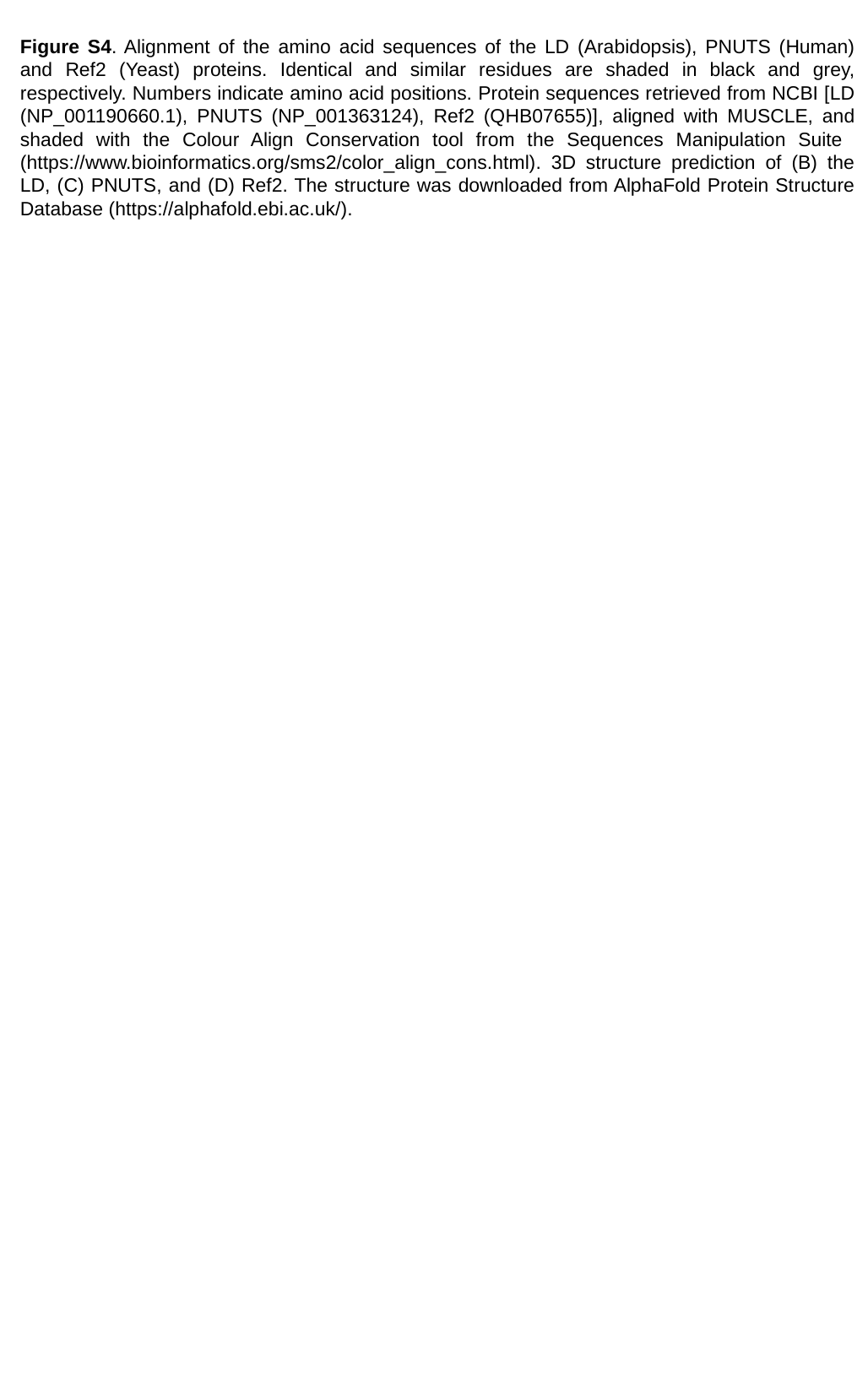

Figure S4. Alignment of the amino acid sequences of the LD (Arabidopsis), PNUTS (Human) and Ref2 (Yeast) proteins. Identical and similar residues are shaded in black and grey, respectively. Numbers indicate amino acid positions. Protein sequences retrieved from NCBI [LD (NP_001190660.1), PNUTS (NP_001363124), Ref2 (QHB07655)], aligned with MUSCLE, and shaded with the Colour Align Conservation tool from the Sequences Manipulation Suite (https://www.bioinformatics.org/sms2/color_align_cons.html). 3D structure prediction of (B) the LD, (C) PNUTS, and (D) Ref2. The structure was downloaded from AlphaFold Protein Structure Database (https://alphafold.ebi.ac.uk/).

### Slide 8
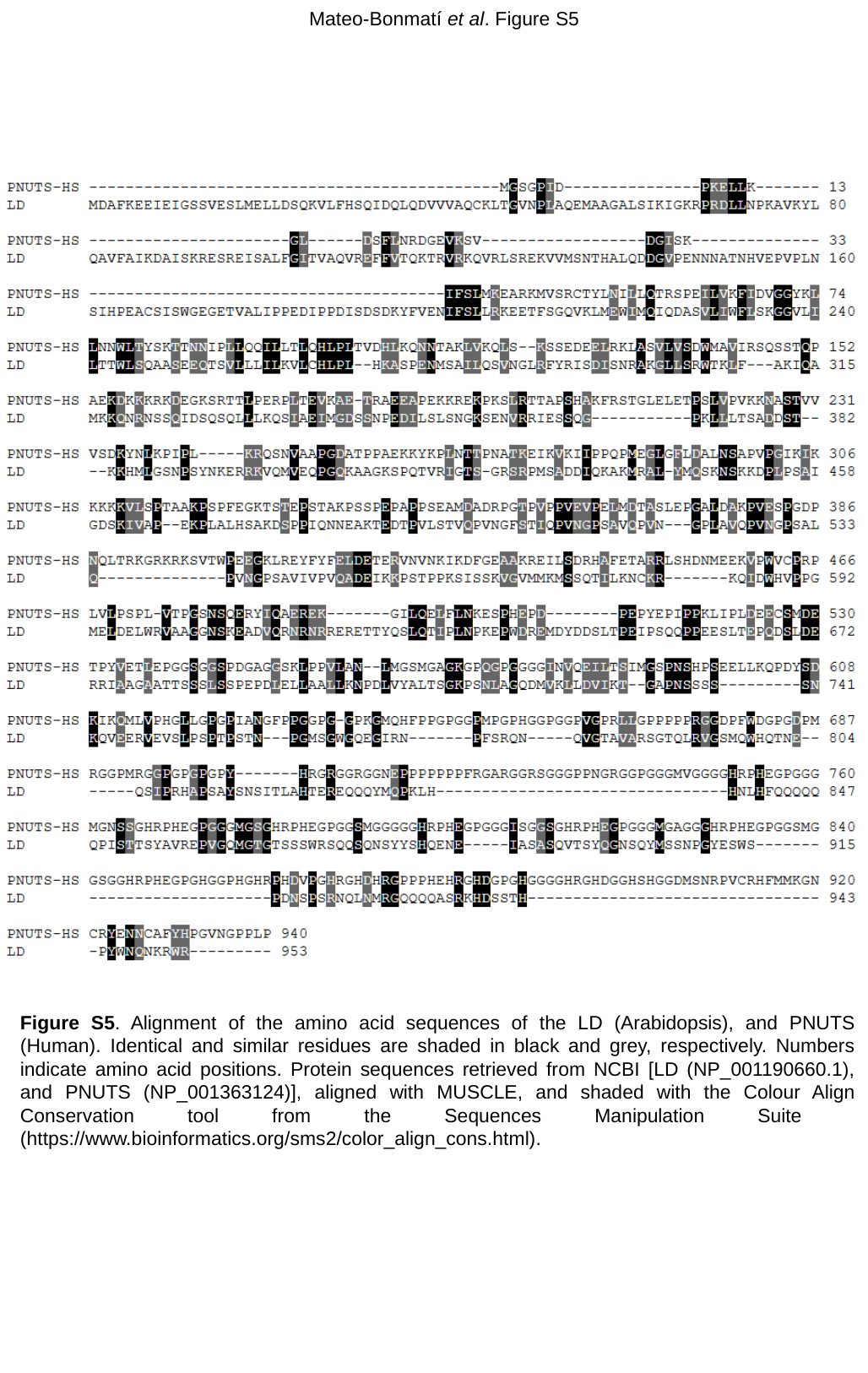

Mateo-Bonmatí et al. Figure S5
Figure S5. Alignment of the amino acid sequences of the LD (Arabidopsis), and PNUTS (Human). Identical and similar residues are shaded in black and grey, respectively. Numbers indicate amino acid positions. Protein sequences retrieved from NCBI [LD (NP_001190660.1), and PNUTS (NP_001363124)], aligned with MUSCLE, and shaded with the Colour Align Conservation tool from the Sequences Manipulation Suite (https://www.bioinformatics.org/sms2/color_align_cons.html).

### Slide 9
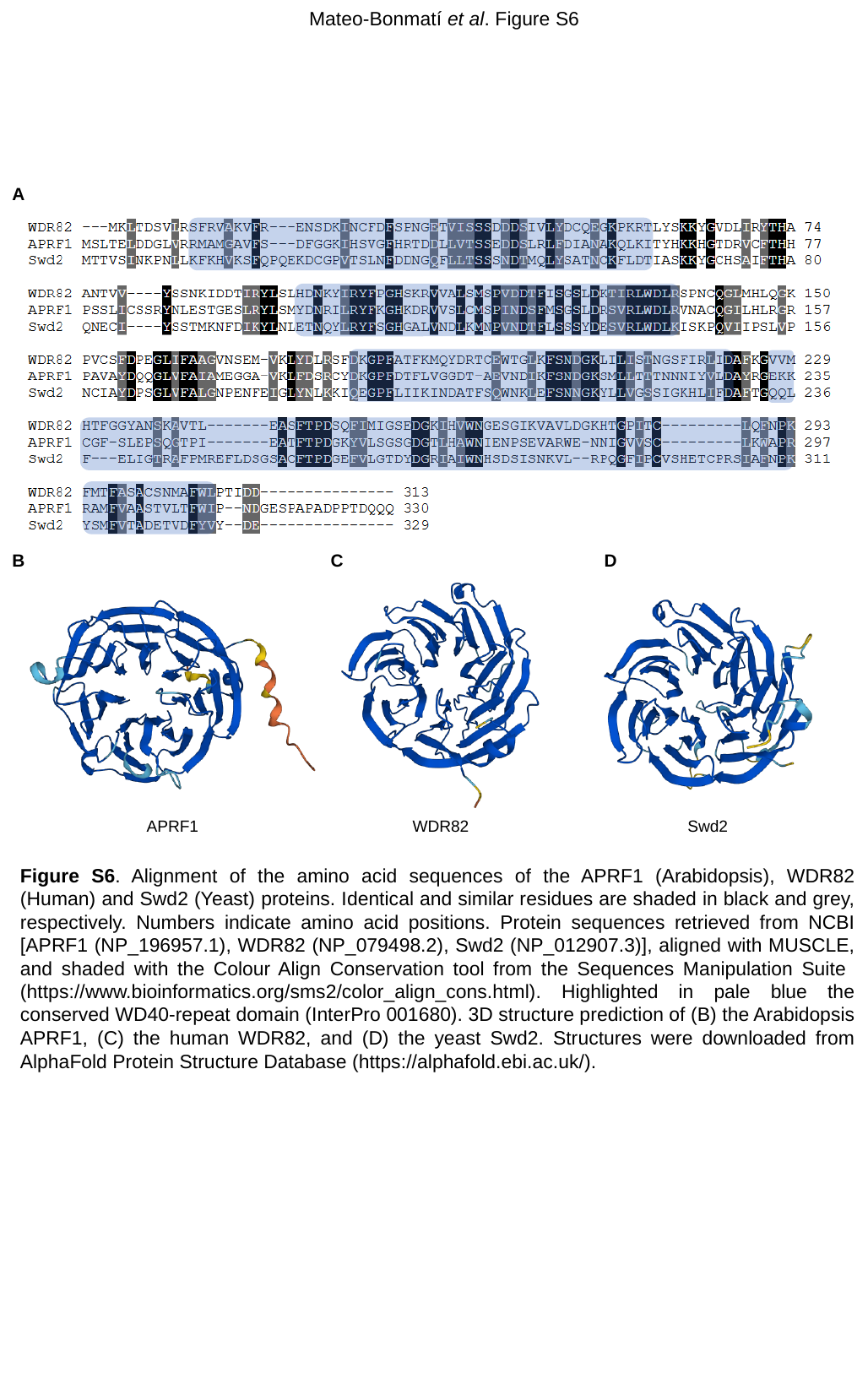

Mateo-Bonmatí et al. Figure S6
A
B
C
D
APRF1
WDR82
Swd2
Figure S6. Alignment of the amino acid sequences of the APRF1 (Arabidopsis), WDR82 (Human) and Swd2 (Yeast) proteins. Identical and similar residues are shaded in black and grey, respectively. Numbers indicate amino acid positions. Protein sequences retrieved from NCBI [APRF1 (NP_196957.1), WDR82 (NP_079498.2), Swd2 (NP_012907.3)], aligned with MUSCLE, and shaded with the Colour Align Conservation tool from the Sequences Manipulation Suite (https://www.bioinformatics.org/sms2/color_align_cons.html). Highlighted in pale blue the conserved WD40-repeat domain (InterPro 001680). 3D structure prediction of (B) the Arabidopsis APRF1, (C) the human WDR82, and (D) the yeast Swd2. Structures were downloaded from AlphaFold Protein Structure Database (https://alphafold.ebi.ac.uk/).

### Slide 10
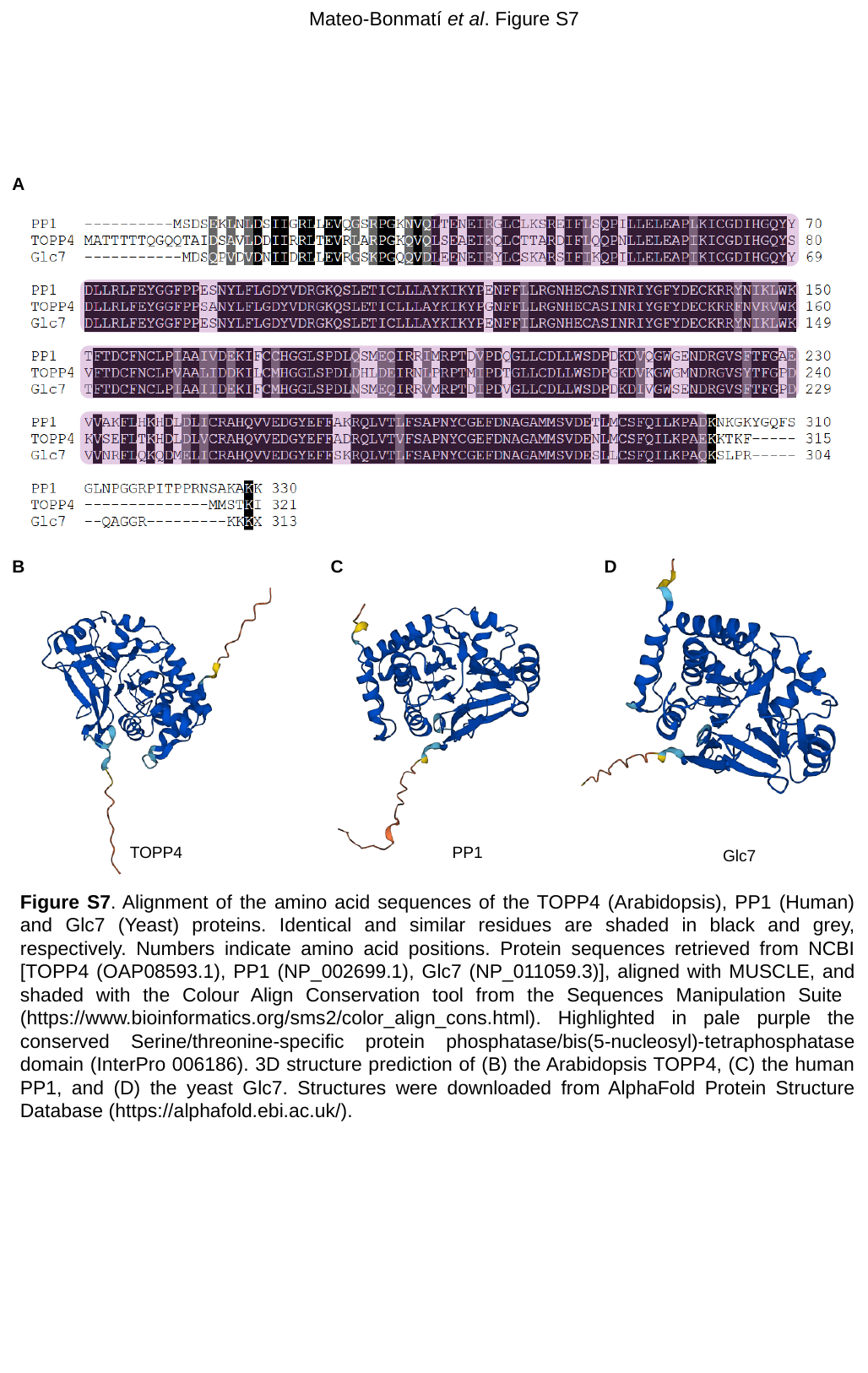

Mateo-Bonmatí et al. Figure S7
A
D
C
B
TOPP4
PP1
Glc7
Figure S7. Alignment of the amino acid sequences of the TOPP4 (Arabidopsis), PP1 (Human) and Glc7 (Yeast) proteins. Identical and similar residues are shaded in black and grey, respectively. Numbers indicate amino acid positions. Protein sequences retrieved from NCBI [TOPP4 (OAP08593.1), PP1 (NP_002699.1), Glc7 (NP_011059.3)], aligned with MUSCLE, and shaded with the Colour Align Conservation tool from the Sequences Manipulation Suite (https://www.bioinformatics.org/sms2/color_align_cons.html). Highlighted in pale purple the conserved Serine/threonine-specific protein phosphatase/bis(5-nucleosyl)-tetraphosphatase domain (InterPro 006186). 3D structure prediction of (B) the Arabidopsis TOPP4, (C) the human PP1, and (D) the yeast Glc7. Structures were downloaded from AlphaFold Protein Structure Database (https://alphafold.ebi.ac.uk/).

### Slide 11
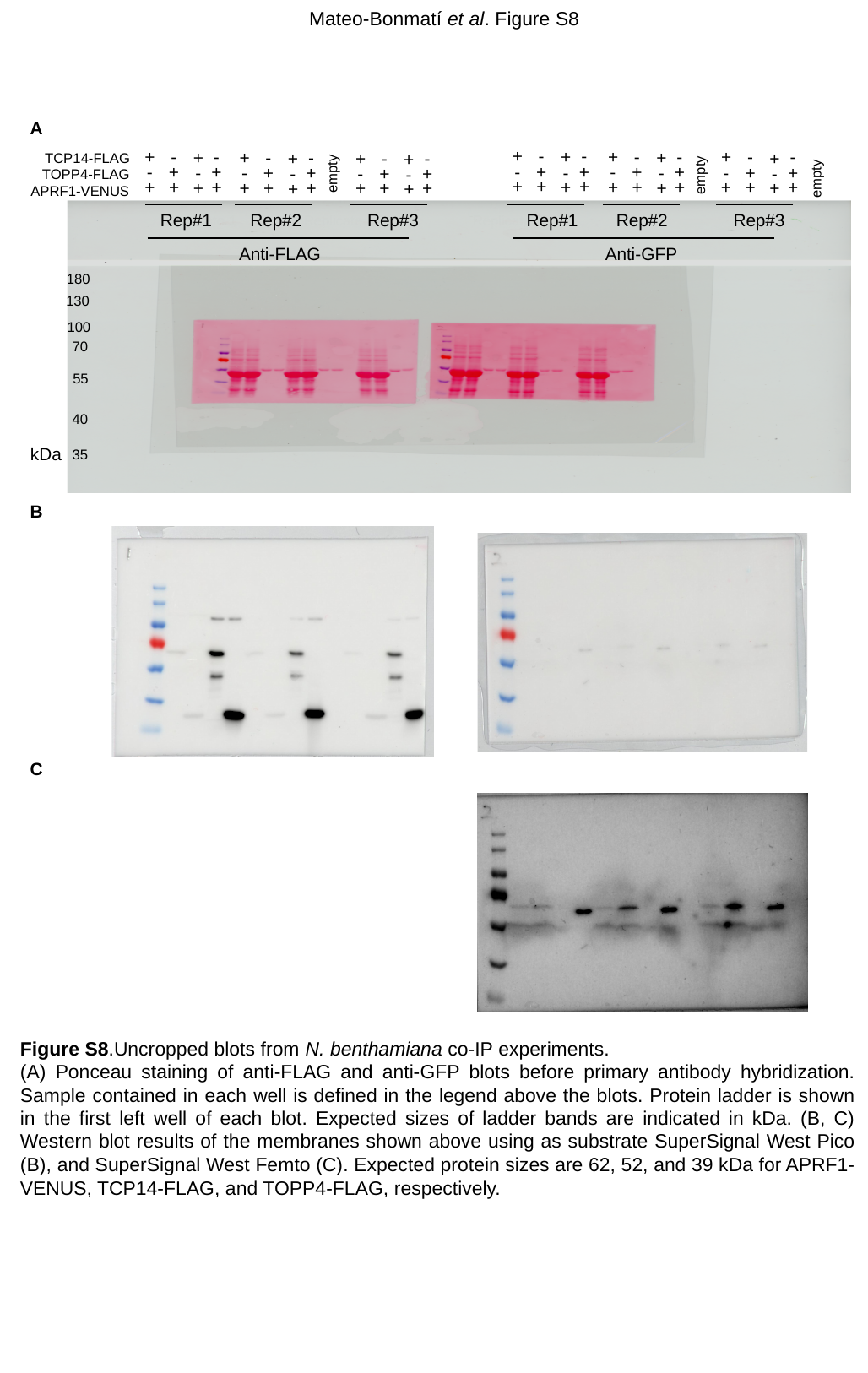

Mateo-Bonmatí et al. Figure S8
A
+
-
-
+
-
-
+
-
-
+
+
-
-
+
-
-
+
+
+
-
-
+
+
+
TCP14-FLAG
-
+
+
-
+
+
-
+
+
-
-
+
+
-
+
+
-
-
-
+
+
-
-
-
empty
TOPP4-FLAG
empty
empty
+
+
+
+
+
+
+
+
+
+
+
+
+
+
+
+
+
+
+
+
+
+
+
+
APRF1-VENUS
Rep#1
Rep#1
Rep#2
Rep#3
Rep#2
Rep#3
Replicate 1
Replicate 3
Replicate 2
Anti-FLAG
Anti-GFP
180
130
100
70
55
40
kDa
35
B
C
Figure S8.Uncropped blots from N. benthamiana co-IP experiments.
(A) Ponceau staining of anti-FLAG and anti-GFP blots before primary antibody hybridization. Sample contained in each well is defined in the legend above the blots. Protein ladder is shown in the first left well of each blot. Expected sizes of ladder bands are indicated in kDa. (B, C) Western blot results of the membranes shown above using as substrate SuperSignal West Pico (B), and SuperSignal West Femto (C). Expected protein sizes are 62, 52, and 39 kDa for APRF1-VENUS, TCP14-FLAG, and TOPP4-FLAG, respectively.

### Slide 12
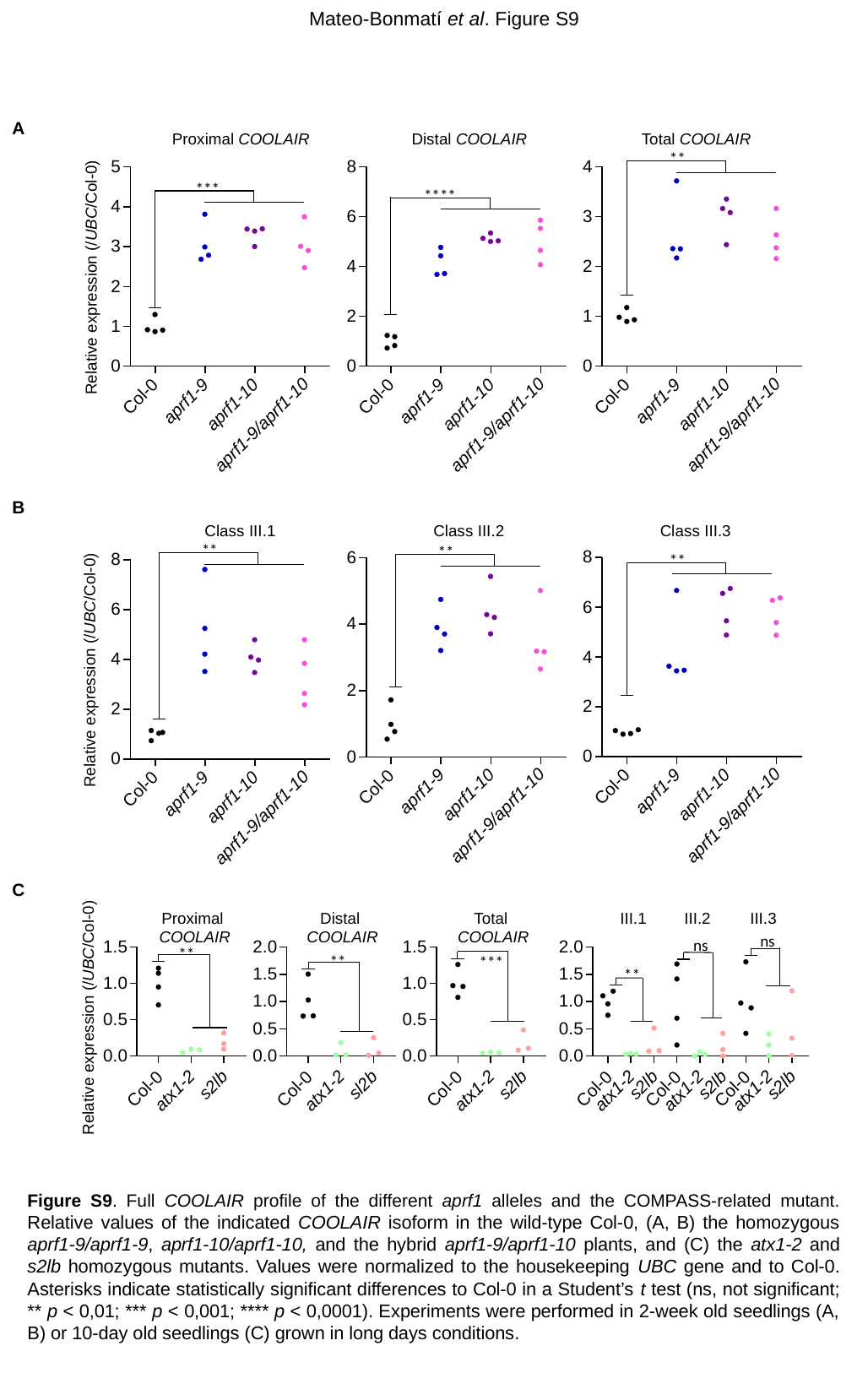

Mateo-Bonmatí et al. Figure S9
A
Proximal COOLAIR
Distal COOLAIR
Total COOLAIR
**
***
****
Relative expression (/UBC/Col-0)
B
Class III.1
Class III.2
Class III.3
**
**
**
Relative expression (/UBC/Col-0)
C
Proximal
COOLAIR
Distal
COOLAIR
Total
COOLAIR
III.1
III.2
III.3
ns
ns
**
**
***
**
Relative expression (/UBC/Col-0)
Figure S9. Full COOLAIR profile of the different aprf1 alleles and the COMPASS-related mutant. Relative values of the indicated COOLAIR isoform in the wild-type Col-0, (A, B) the homozygous aprf1-9/aprf1-9, aprf1-10/aprf1-10, and the hybrid aprf1-9/aprf1-10 plants, and (C) the atx1-2 and s2lb homozygous mutants. Values were normalized to the housekeeping UBC gene and to Col-0. Asterisks indicate statistically significant differences to Col-0 in a Student’s t test (ns, not significant; ** p < 0,01; *** p < 0,001; **** p < 0,0001). Experiments were performed in 2-week old seedlings (A, B) or 10-day old seedlings (C) grown in long days conditions.

### Slide 13
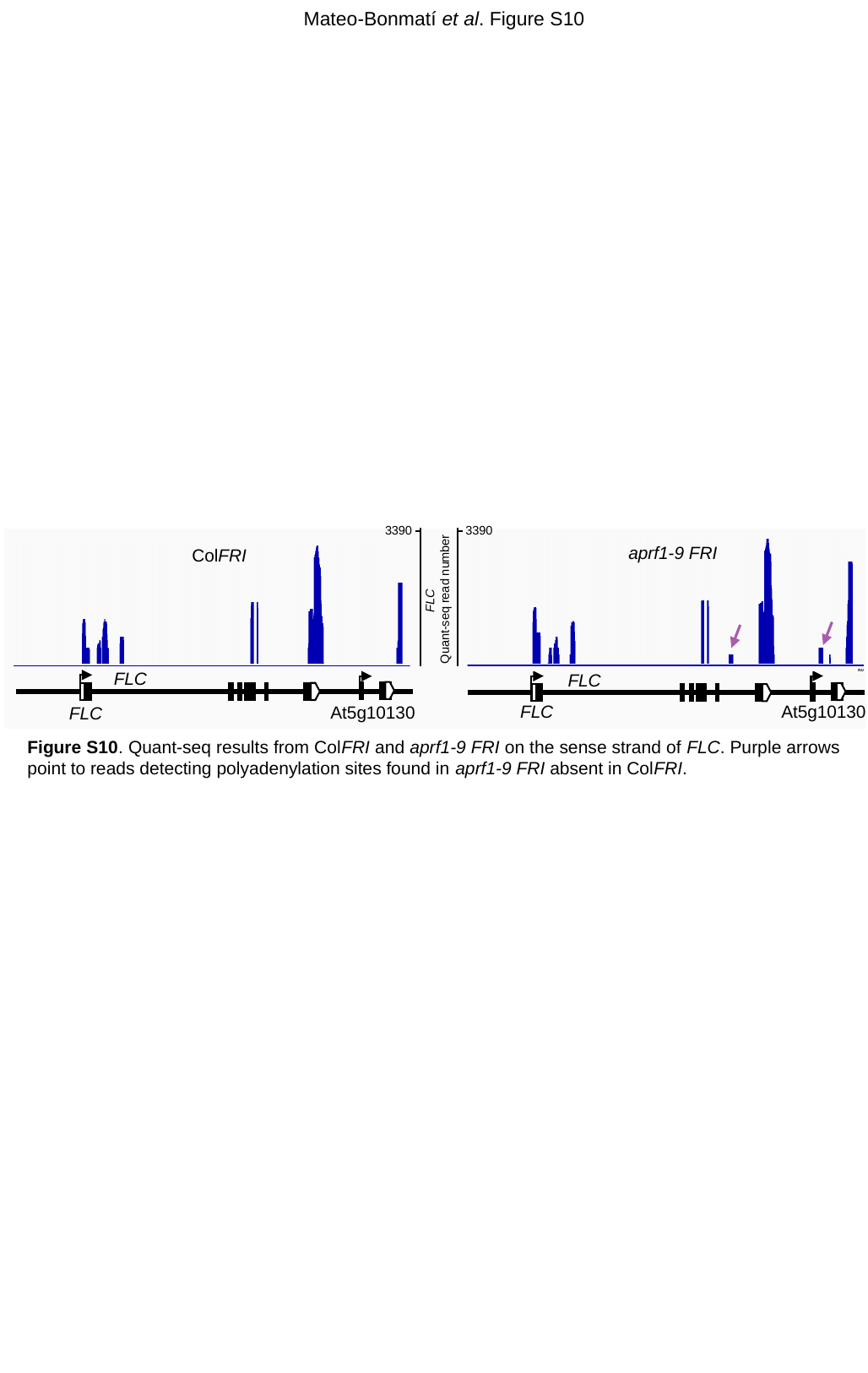

Mateo-Bonmatí et al. Figure S10
3390
3390
aprf1-9 FRI
ColFRI
FLC
Quant-seq read number
FLC
FLC
At5g10130
FLC
At5g10130
FLC
Figure S10. Quant-seq results from ColFRI and aprf1-9 FRI on the sense strand of FLC. Purple arrows point to reads detecting polyadenylation sites found in aprf1-9 FRI absent in ColFRI.

### Slide 14
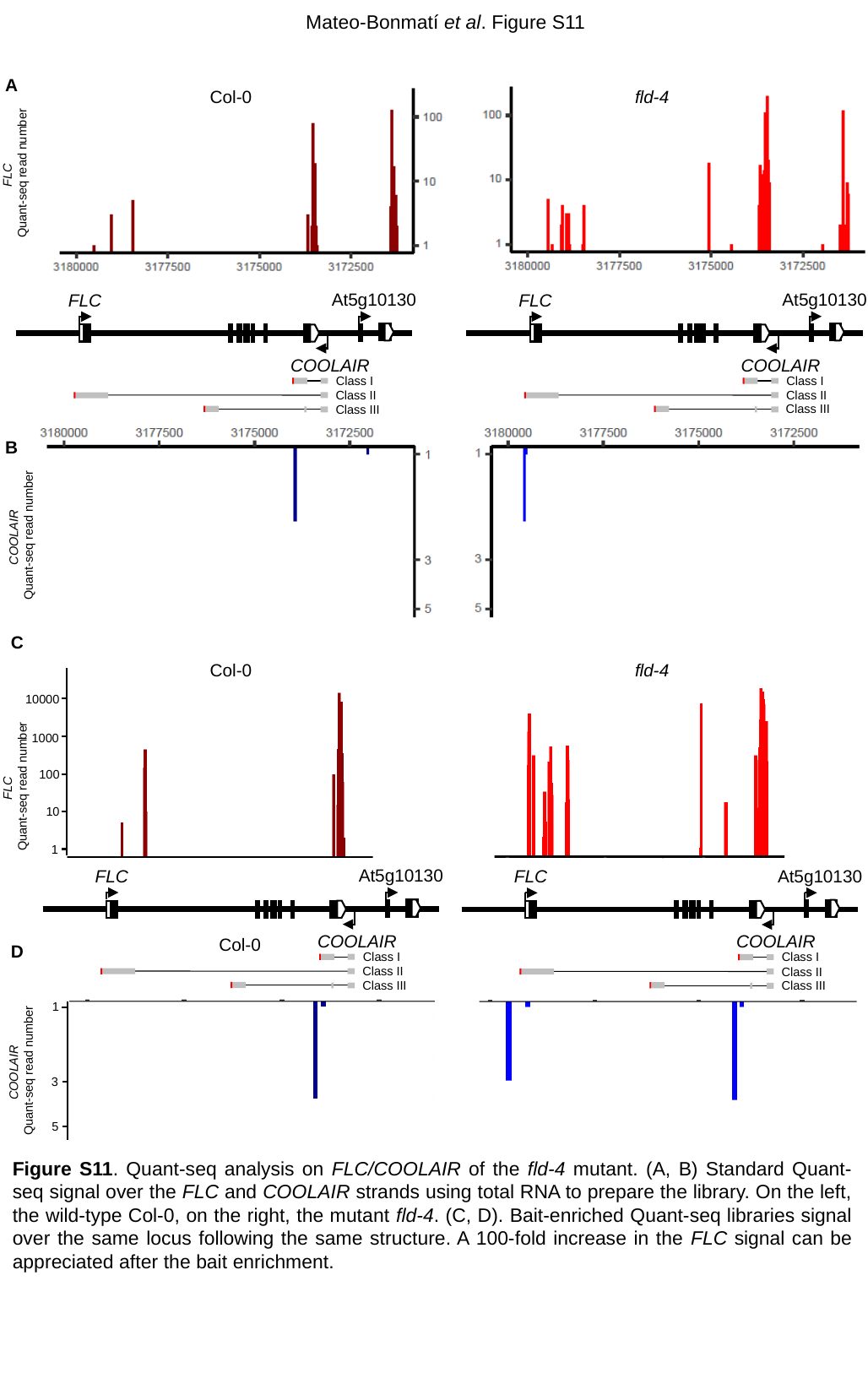

Mateo-Bonmatí et al. Figure S11
A
Col-0
fld-4
FLC
Quant-seq read number
At5g10130
At5g10130
FLC
COOLAIR
FLC
COOLAIR
Class I
Class I
Class II
Class II
B
COOLAIR
Quant-seq read number
Class III
Class III
C
Col-0
fld-4
10000
1000
100
FLC
Quant-seq read number
10
fld-4
1
At5g10130
At5g10130
FLC
COOLAIR
FLC
COOLAIR
Col-0
D
Class I
Class I
Class II
Class II
Class III
Class III
1
COOLAIR
Quant-seq read number
3
5
Figure S11. Quant-seq analysis on FLC/COOLAIR of the fld-4 mutant. (A, B) Standard Quant-seq signal over the FLC and COOLAIR strands using total RNA to prepare the library. On the left, the wild-type Col-0, on the right, the mutant fld-4. (C, D). Bait-enriched Quant-seq libraries signal over the same locus following the same structure. A 100-fold increase in the FLC signal can be appreciated after the bait enrichment.

### Slide 15
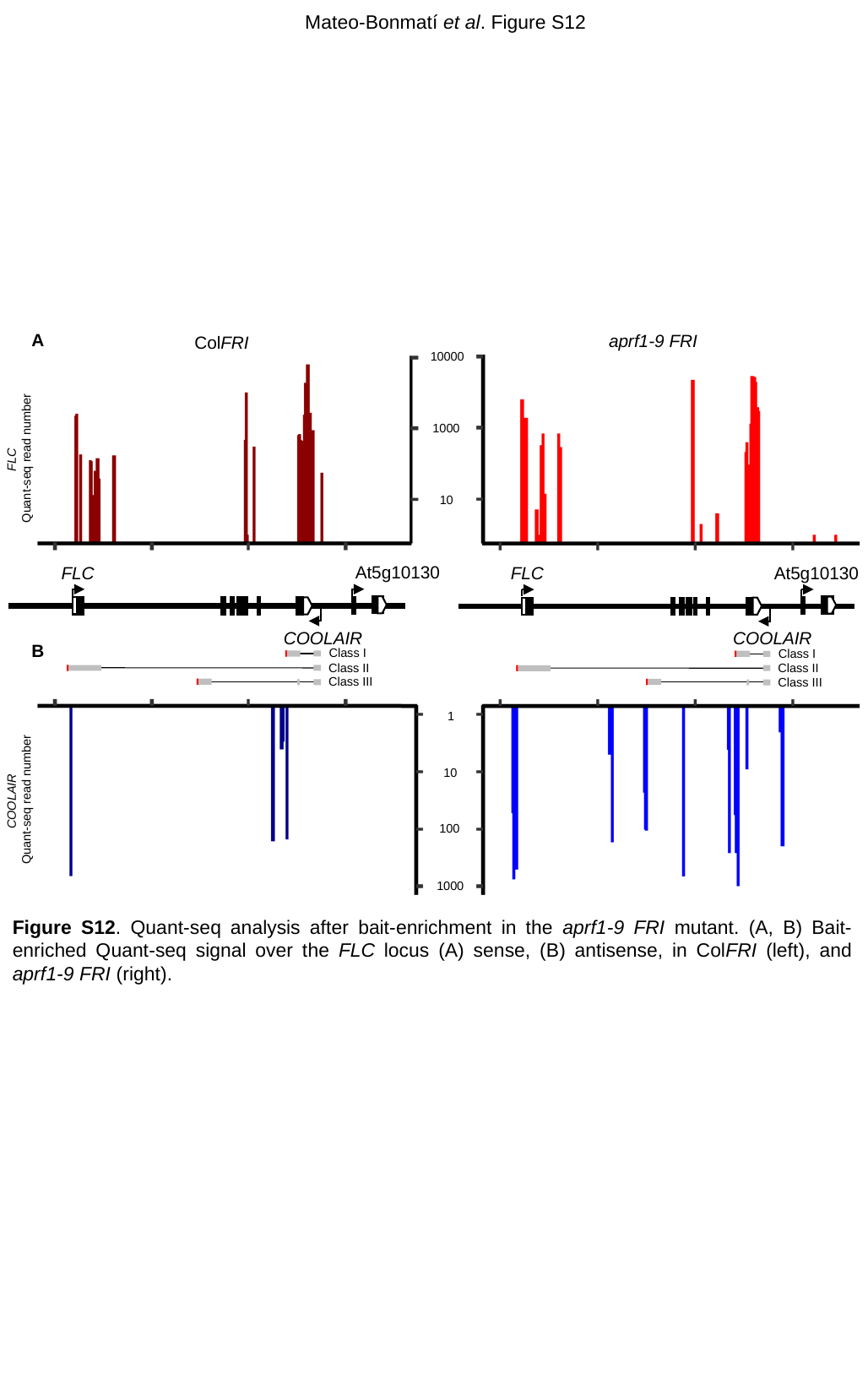

Mateo-Bonmatí et al. Figure S12
A
aprf1-9 FRI
ColFRI
At5g10130
At5g10130
FLC
COOLAIR
FLC
COOLAIR
Class I
Class I
Class II
Class II
Class III
Class III
10000
1000
FLC
Quant-seq read number
10
B
1
10
COOLAIR
Quant-seq read number
100
1000
Figure S12. Quant-seq analysis after bait-enrichment in the aprf1-9 FRI mutant. (A, B) Bait-enriched Quant-seq signal over the FLC locus (A) sense, (B) antisense, in ColFRI (left), and aprf1-9 FRI (right).

### Slide 16
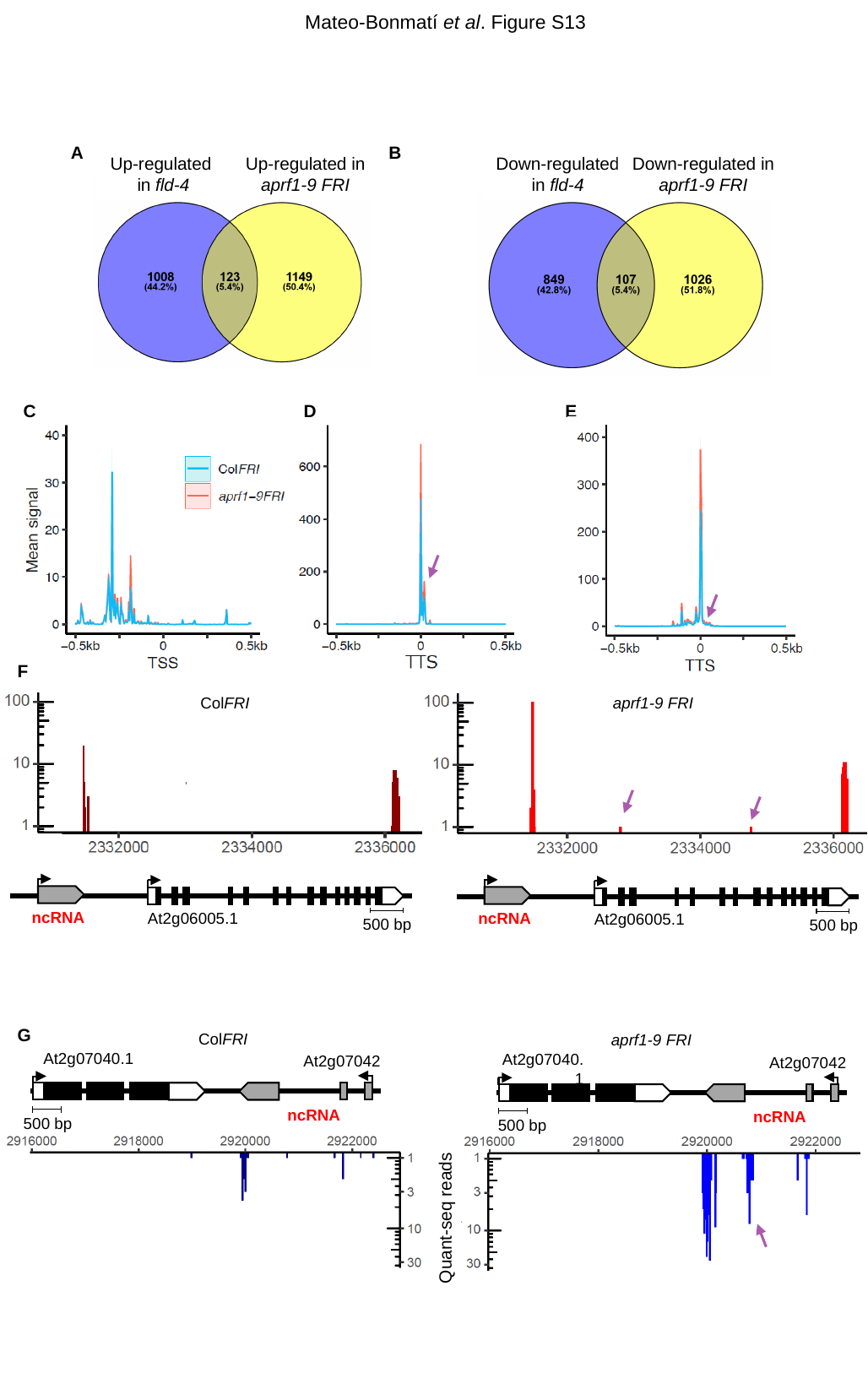

Mateo-Bonmatí et al. Figure S13
A
B
Up-regulated
in fld-4
Up-regulated in
aprf1-9 FRI
Down-regulated
in fld-4
Down-regulated in
aprf1-9 FRI
E
C
D
F
ColFRI
aprf1-9 FRI
ncRNA
At2g06005.1
500 bp
ncRNA
At2g06005.1
500 bp
G
ColFRI
aprf1-9 FRI
At2g07040.1
At2g07042
ncRNA
500 bp
At2g07040.1
At2g07042
ncRNA
500 bp
Quant-seq reads

### Slide 17
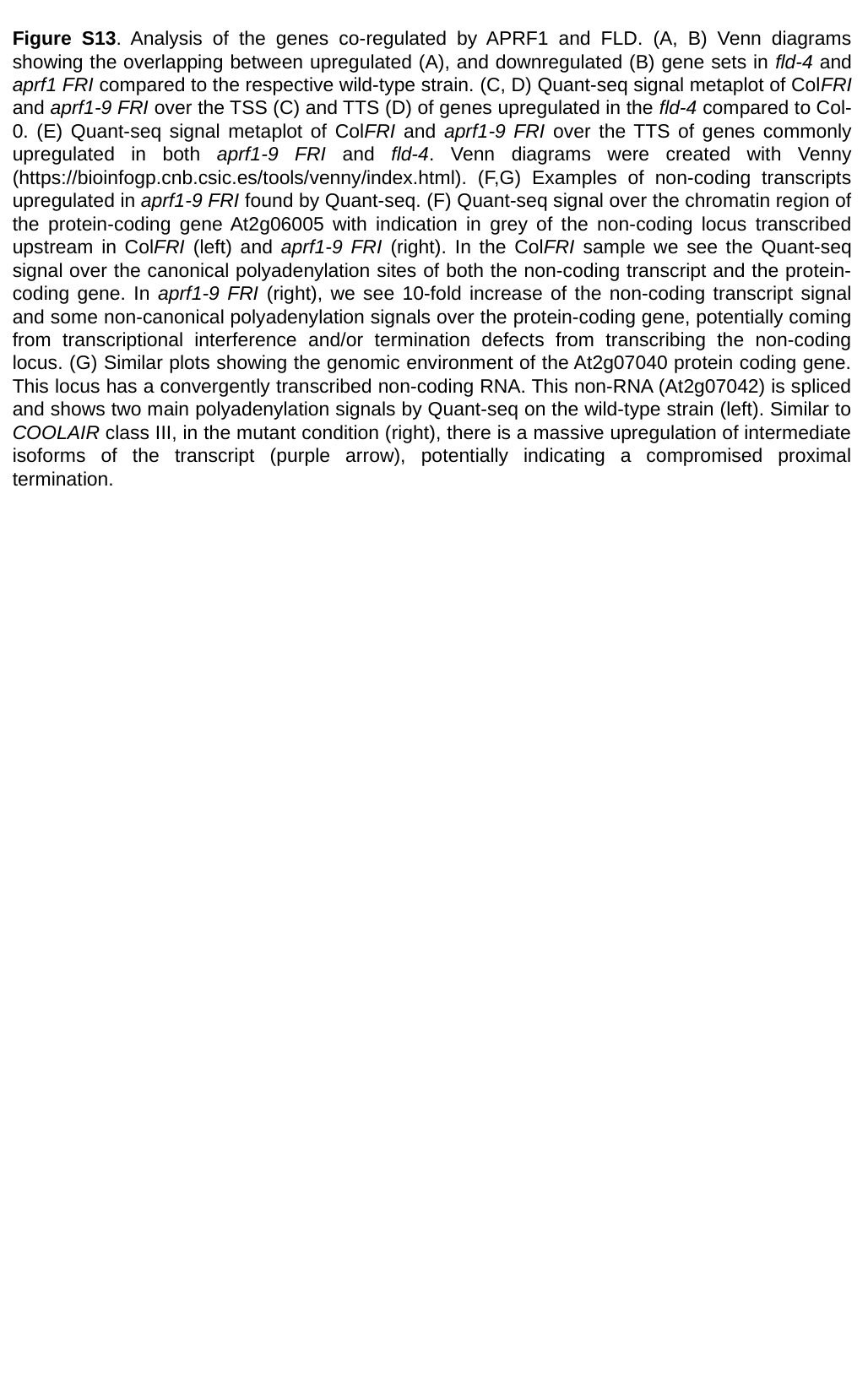

Figure S13. Analysis of the genes co-regulated by APRF1 and FLD. (A, B) Venn diagrams showing the overlapping between upregulated (A), and downregulated (B) gene sets in fld-4 and aprf1 FRI compared to the respective wild-type strain. (C, D) Quant-seq signal metaplot of ColFRI and aprf1-9 FRI over the TSS (C) and TTS (D) of genes upregulated in the fld-4 compared to Col-0. (E) Quant-seq signal metaplot of ColFRI and aprf1-9 FRI over the TTS of genes commonly upregulated in both aprf1-9 FRI and fld-4. Venn diagrams were created with Venny (https://bioinfogp.cnb.csic.es/tools/venny/index.html). (F,G) Examples of non-coding transcripts upregulated in aprf1-9 FRI found by Quant-seq. (F) Quant-seq signal over the chromatin region of the protein-coding gene At2g06005 with indication in grey of the non-coding locus transcribed upstream in ColFRI (left) and aprf1-9 FRI (right). In the ColFRI sample we see the Quant-seq signal over the canonical polyadenylation sites of both the non-coding transcript and the protein-coding gene. In aprf1-9 FRI (right), we see 10-fold increase of the non-coding transcript signal and some non-canonical polyadenylation signals over the protein-coding gene, potentially coming from transcriptional interference and/or termination defects from transcribing the non-coding locus. (G) Similar plots showing the genomic environment of the At2g07040 protein coding gene. This locus has a convergently transcribed non-coding RNA. This non-RNA (At2g07042) is spliced and shows two main polyadenylation signals by Quant-seq on the wild-type strain (left). Similar to COOLAIR class III, in the mutant condition (right), there is a massive upregulation of intermediate isoforms of the transcript (purple arrow), potentially indicating a compromised proximal termination.

### Slide 18
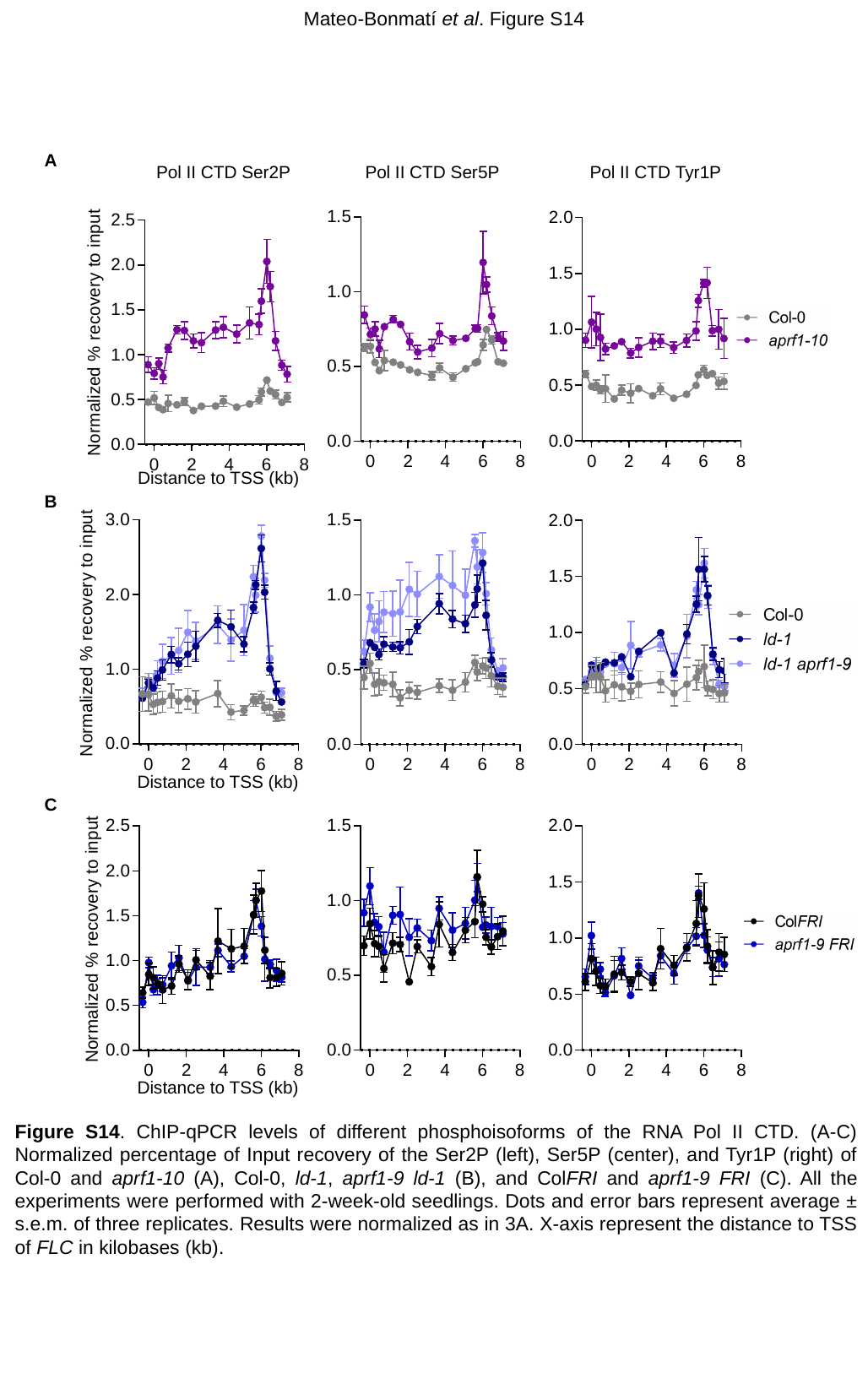

Mateo-Bonmatí et al. Figure S14
A
Pol II CTD Ser2P
Pol II CTD Ser5P
Pol II CTD Tyr1P
Distance to TSS (kb)
B
Distance to TSS (kb)
C
Distance to TSS (kb)
Figure S14. ChIP-qPCR levels of different phosphoisoforms of the RNA Pol II CTD. (A-C) Normalized percentage of Input recovery of the Ser2P (left), Ser5P (center), and Tyr1P (right) of Col-0 and aprf1-10 (A), Col-0, ld-1, aprf1-9 ld-1 (B), and ColFRI and aprf1-9 FRI (C). All the experiments were performed with 2-week-old seedlings. Dots and error bars represent average ± s.e.m. of three replicates. Results were normalized as in 3A. X-axis represent the distance to TSS of FLC in kilobases (kb).

### Slide 19
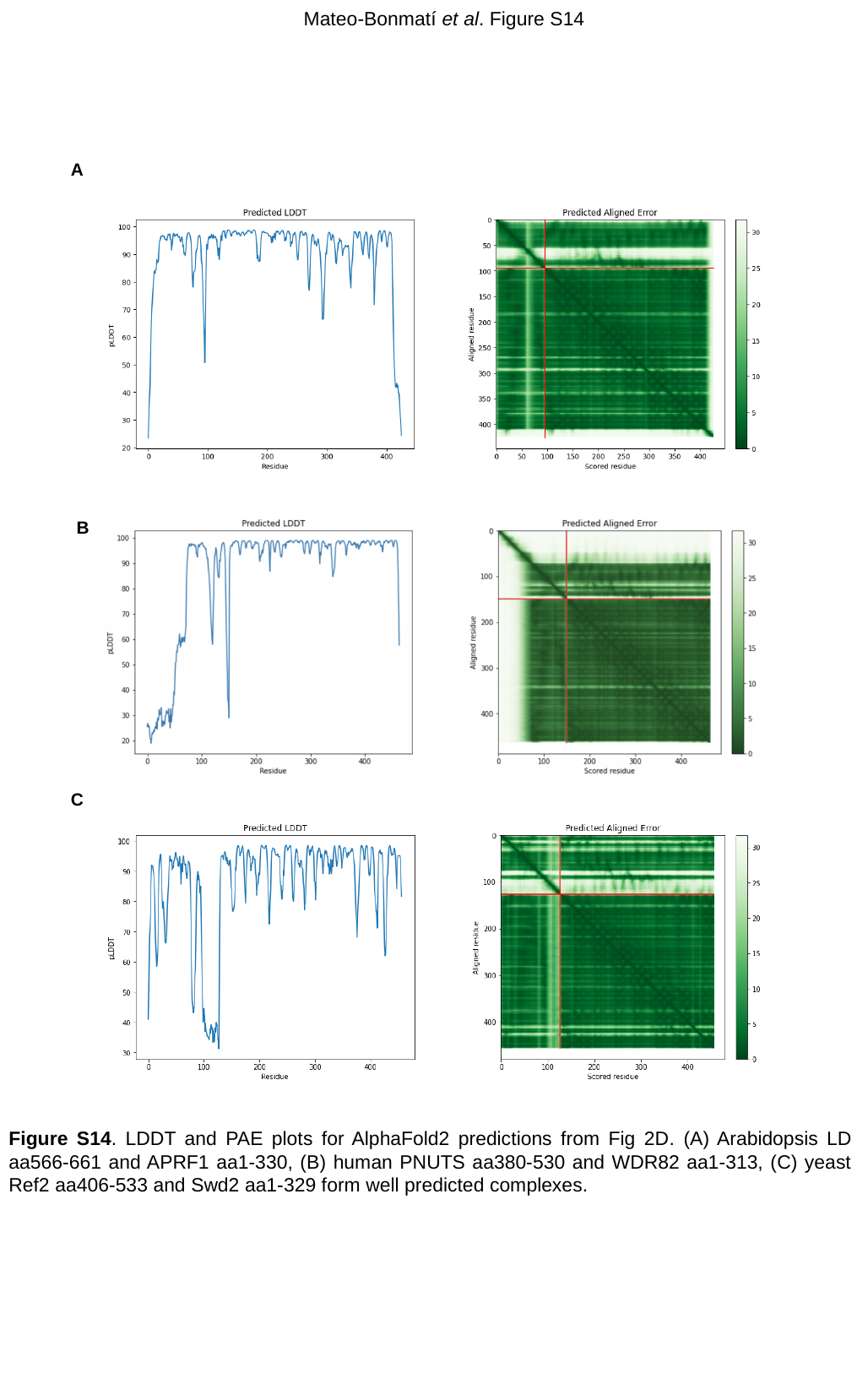

Mateo-Bonmatí et al. Figure S14
A
B
C
Figure S14. LDDT and PAE plots for AlphaFold2 predictions from Fig 2D. (A) Arabidopsis LD aa566-661 and APRF1 aa1-330, (B) human PNUTS aa380-530 and WDR82 aa1-313, (C) yeast Ref2 aa406-533 and Swd2 aa1-329 form well predicted complexes.
