## Supplementary material for "A CPF-like phosphatase module links transcription termination to chromatin silencing": Table S4

| **Table S4**. Primer sets used in this work | | | |
| --- | --- | --- | --- |
| Purpose | Oligonucleotide name | Oligonucleotide sequences (5’ → 3’) | |
|  |  | Forward primer (F) | Reverse primer (R) |
| Genotyping | WiscDsLox489-492K11 | GTTTTTCGAGCAAAGGGAAAG | ACAGGGTACACCCATAGAGGC |
|  | LB_WiscDs |  | TCCTCGAGTTTCTCCATAATAATGT |
|  | dCAPS_aprf1-10 (MaeII) |  |  |
|  | dCAPS_fca-9 (Sty1) | TGTTGAGATGGTGAAACTGTG | TCTTTGGCTCAGCAAACC |
|  | fld-4 | AGAAACCTGCCTGAATGTC | AGCTAGGCAACTGATGAG |
|  | LBb1.3 |  | ATTTTGCCGATTTCGGAAC |
|  | ld-1 | TGGTACAGGAGGATTGAATCG | CTCTGAACATCAGCCTCCTTG |
|  | LB3 |  | TAGCATCTGAATTTCATAACCAATCTC |
|  | FRI | AGATTTGCTGGATTTGATAAGG | CTTGATGTTGGTCGATGATG |
|  | 35S:FCA | GCCACTTGCTCTCTCCTCAC | TTTCAATTCTCTCTGCCTCTCA |
|  | s2lb | TGACAGCTAGCGACGATGAC | TGGTTGTAAGCAGCATGAGC |
|  | LB (s2lb) |  | CCGGATCGTATCGGTTTTCG |
| Cloning | sgRNA_APRF1 | AATGGGAGGGAAGATACATTCGGT | AAACACCGAATGTATCTTCCCTCC |
|  | pKIR | AGAAGAGAAGCAGGCCCATT | AATGGATTGAGCCAAAGAGC |
|  | APRF1-mVENUS | GGCCAGTGCCAAGCTAAGGATCGAAATTCCCCCGAG | CGGCAGCAGAAAGCTTCTGTTGTTGATCGGTAGGAGG |
|  | TOPP4-3xFLAG | CCGGGTACCGAGCTCGAATTCCAAAAACGCAATTTAACATTTTTCCCATCA | CCTTATAGTCAATCTTTGTGGACATCATGAACTTGGT |
| RT | FLC spliced |  | TTTGTCCAGCAGGTGACATC |
|  | FLC unspliced |  | CTTTGTAATCAAAGGTGGAGAGC |
|  | UBC |  | TTGTGCCATTGAATTGAACCC |
|  | PP2A |  | CCAAGCATGGCCGTATCATGT |
|  | Total COOLAIR | TGCATCGAGATCTTGAGTGTATGT |  |
|  | Proximal COOLAIR | TGGTTGTTATTTGGTGGTGTG |  |
| RT | Distal COOLAIR | GCCCGACGAAGAAAAAGTAG |  |
| (continuation) | Class III COOLAIR | GAAACAATCTGGACAGTAGAGGC |  |
| qPCR | FLC spliced | AGCCAAGAAGACCGAACTCA | TTTGTCCAGCAGGTGACATC |
|  | FLC unspliced | CGCAATTTTCATAGCCCTTG | CTTTGTAATCAAAGGTGGAGAGC |
|  | Proximal COOLAIR | CCTGCTGGACAAATCTCCGA | TCACACGAATAAGGTGGCTAATTAAG |
|  | Distal COOLAIR | GTATCTCCGGCGACTTGAAC | GGATGCGTCACAGAGAACAG |
|  | Total COOLAIR | TGCATCGAGATCTTGAGTGTATGT | ACGTCCCTGTTGCAAAATAAGC |
|  | COOLAIR class III.1 | AGTAGAGGCTTATGTTTAGGGTTCT | TCTCACACGAATAAGATTGAAAATGAC |
|  | COOLAIR class III.2 | AGGCTTATGTTTAGGGTTCTTATGTAC | TCCATCTGTACGATAATCATAGATTGA |
|  | COOLAIR class III.3 | GAAACAATCTGGACAGTAGAGGC | TGTCCAGCAGATTGAAAATGACA |
|  | PP2A | ACTGCATCTAAAGACAGAGTTCC | CCAAGCATGGCCGTATCATGT |
|  | UBC | CTGCGACTCAGGGAATCTTCTAA | TTGTGCCATTGAATTGAACCC |
| ChIP | FLC_-2285 | ATCCAGAAAAGGGCAAGGAG | CGAATCGATTGGGTGAATG |
|  | FLC_-1788 | GGATTGATGTGGGGCACTAT | AGTCATGGGTAGGGCATGTG |
|  | FLC_-1555 | TGGAGGGAACAACCTAATGC | TCATTGGACCAAACCAAACC |
|  | FLC_-321 | ACTATGTAGGCACGACTTTGGTAAC | TGCAGAAAGAACCTCCACTCTAC |
|  | FLC_5 | GCCCGACGAAGAAAAAGTAG | TCCTCAGGTTTGGGTTCAAG |
|  | FLC_246 | CGACAAGTCACCTTCTCCAAA | AGGGGGAACAAATGAAAACC |
|  | FLC_473 | GGCGGATCTCTTGTTGTTTC | CTTCTTCACGACATTGTTCTTCC |
|  | FLC_741 | CGTGCTCGATGTTGTTGAGT | TCCCGTAAGTGCATTGCATA |
|  | FLC_1212 | CCTTTTGCTGTACATAAACTGGTC | CCAAACTTCTTGATCCTTTTTACC |
|  | FLC_1613 | TTGACAATCCACAACCTCAATC | TCAATTTCCTAGAGGCACCAA |
|  | FLC_2094 | AGCCTTTTAGAACGTGGAACC | TCTTCCATAGAAGGAAGCGACT |
|  | FLC_2523 | AGTTTGGCTTCCTCATACTTATGG | CAATGAACCTTGAGGACAAGG |
|  | FLC_3276 | GGGGCTGCGTTTACATTTTA | GTGATAGCGCTGGCTTTGAT |
|  | FLC_3699 | TGAAATGTTACGAATACTAGCGTGT | GGATCAAAACTACTAGCTAACCCTTG |
|  | FLC_4406 | AGAACAACCGTGCTGCTTTT | TGTGTGCAAGCTCGTTAAGC |
|  | FLC_5090 | CCGGTTGTTGGACATAACTAGG | CCAAACCCAGACTTAACCAGAC |
|  | FLC_5599 | TGGTTGTTATTTGGTGGTGTG | ATCTCCATCTCAGCTTCTGCTC |
|  | FLC_5715 | CCTGCTGGACAAATCTCCGA | GGATTTTGATTTCAACCGCCGA |
| ChIP | FLC_6013 | CGTGTGAGAATTGCATCGAG | AAAAACGCGCAGAGAGAGAG |
| (continuation) | FLC_6189 | TCCTAAACGCGTATGGTTGG | CCTTCATGGATGACGGAACT |
|  | FLC_6480 | TCCAGACGCCATTGTCATTA | AGAGTGCATTTTAACACTGACGA |
|  | FLC_6810 | TTGTAAAGTCCGATGGAGACG | ACTCGGCGAGAAAGTTTGTG |
|  | FLC_7091 | CATCGCTGTGTTATGCGTTT | GGCAAGTGTCGCAGTAAACA |
|  | STM (for H3K27me3 normalization) | GCCCATCATGACATCACATC | GGGAACTACTTTGTTGGTGGTG |
|  | ACT-914 (for RNA Pol II ChIP normalization) | TGGGTCTCATATAGAACACTCACAAAGGT | GACCAAAACCCGAATAGGAGCAAGA |
|  | ACT_+122 (for H3K36me3 normalization) | CGTTTCGCTTTCCTTAGTGTTAGCT | AGCGAACGGATCTAGAGACTCACCTTG |
|  | ACT+939 (for H3K4me1 normalization) | TGCCCCGAGAGCAGTGTTCC | TGGACTGAGCTTCATCACCAACG |
